# Intra- and inter-species interactions drive early phases of invasion in mice gut microbiota

**DOI:** 10.1101/2022.12.30.522336

**Authors:** Melis Gencel, Gisela Marrero Cofino, Cang Hui, Zahra Sahaf, Louis Gauthier, Derek Tsang, Dana Philpott, Sheela Ramathan, Alfredo Menendez, Shimon Bershtein, Adrian W.R. Serohijos

## Abstract

The stability and dynamics of ecological communities are dictated by interaction networks typically quantified at the level of species.^1–10^ But how such networks are influenced by intra-species variation (ISV) is poorly understood.^11–14^ Here, we use ~500,000 chromosomal barcodes to track high-resolution intra-species clonal lineages of *Escherichia coli* invading mice gut with the increasing complexity of gut microbiome: germ-free, antibiotic-perturbed, and innate microbiota. By co-clustering the dynamics of intra-species clonal lineages and those of gut bacteria from 16S rRNA profiling, we show the emergence of complex time-dependent interactions between *E. coli* clones and resident gut bacteria. With a new approach, dynamic covariance mapping (DCM), we differentiate three phases of invasion in susceptible communities: 1) initial loss of community stability as *E. coli* enters; 2) recolonization of some gut bacteria; and 3) recovery of stability with *E. coli* coexisting with resident bacteria in a quasi-steady state. Comparison of the dynamics, stability and fitness from experimental replicates and different cohorts suggest that phase 1 is driven by mutations in *E. coli* before colonization, while phase 3 is by *de novo* mutations. Our results highlight the transient nature of interaction networks in microbiomes driven by the persistent coupling of ecological and evolutionary dynamics.

**One-Sentence Summary:** High-resolution lineage tracking and dynamic covariance mapping (DCM) define three distinct phases during early gut microbiome invasion.

## Introduction

How an ecological network responds to perturbation, such as an invasion, depends on the structure of the community interaction matrix^1,2^ that quantifies pairwise effects of each species on other’s population growth. Due to limited experimental resolution, this matrix is typically described at the level of species or higher-level taxonomic groupings^3–10,15,16^. This core concept of ecology has been scrutinized in diverse systems of plants, animals, and microbes under both natural and lab conditions^7,10,17–21^. However, a species rarely exists as a homogenous population due to spatial partitioning and genomic variation from pre-existing or *de novo* mutations^22^. Despite the prevalence of intra-species variation (ISV) in nature, how the ISV of an invading species affects the community matrix is not fully known. Indeed, the role of ISV on community composition and stability has rarely been tested experimentally^11–13^. Theoretical studies have also yielded contradictory results regarding the role of intraspecific variation on species coexistence^14^.

Here, we experimentally determine the impact of intraspecific variation on community dynamics during *E. coli* invasion in the mouse gut microbiome. The mouse gut microbiome, depending on host phenotype and genetics, consists of ~10^12^ individual bacterial cells, which can be partitioned into ~10^3^ species^23^. The relatively high mutation rate, large population size, and frequency-dependent selection could lead to ~10^5^ clones existing in a population of a single bacterial species. Some of these clones present at very low frequency could nonetheless provide rapid adaptation^24,25^. Numerous whole-genome sequencing studies with barcoding technologies highlight the diversity of intra-species clonal lineages^24–29^, although limited to a few thousand cells of a single species or hundreds of cells in species-rich consortia. This shallow coverage of intra-species lineage dynamics is yet to determine the extent of coupling between ecological and evolutionary processes^30,31^. Instead, deciphering the effect of intra-species clonal variation on community dynamics requires the ability to track intraspecific clonal lineages at very high resolutions.

We used high-density chromosomal barcoding to track the clonal lineages of *E. coli* that were introduced into mice with different degrees of complexity in their resident bacterial community. First, this enabled us to define the ISV dynamics at a frequency as low as 1 in every ~10^7^ cells of *E. coli*. Second, the ISV dynamics enabled us to define the dominant and persistent *E. coli* clonal lineages (“clones”), including those present at low frequencies. Third, by correlating this ISV dynamics with the composition of the resident bacteria from 16S profiling, we found that specific clonal lineages interact with specific resident bacterial families. Notably, such interactions were reproducible across mice colonization replicates that were susceptible to *E. coli* invasion. Fourth, we expand the definition of community matrix interactions, traditionally defined between species or families, to include both intra-species and between-species interactions. This resulted in a new approach, which we call dynamic covariance mapping (DCM), which could quantify the time-dependence of the community interaction matrix and its effect on community stability. Fifth, using DCM, we were able to define distinct temporal phases during colonization, which uniquely arise due to the specific interaction between *E. coli* clones and certain families of the resident microbiota.

## Results

### High-resolution clonal tracking during gut bacterial community invasion

We previously used the Tn7 transposon machinery to introduce ~500,000 distinct chromosomal DNA barcodes into a population of ~10^8^ *E. coli* cells^24^. Since the barcodes are transmitted from parent to daughter cells, this allowed the tracking of the clonal lineage dynamics of *E. coli* at a resolution of ~1/10^6^ cells as the population adapted to antibiotic resistance during *in vitro* lab evolution^24^. Such high-resolution chromosomal barcoding techniques have been used for single-species analysis but never in a complex and species-rich ecological community^32^. We used barcodes to simultaneously track high-resolution clonal lineage dynamics of an *E. coli* population colonizing mouse guts (Fig. 1a).

**Figure 1.**
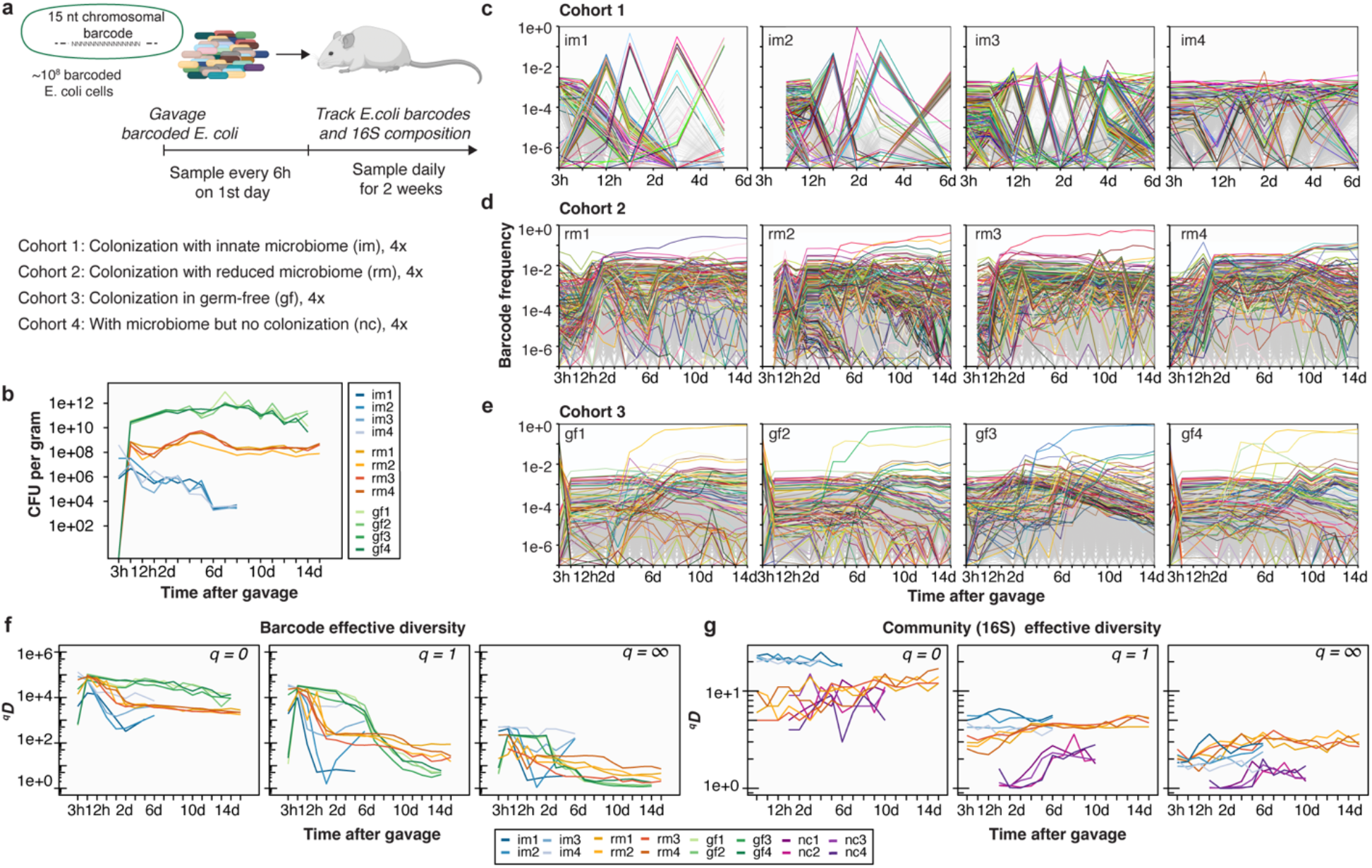
Intra-species population dynamics during gut colonization. **a,** Population of ~10^8^ *E. coli* cells with ~5×10^5^ unique chromosomal barcodes is introduced into mice with innate microbiota (cohort 1) and mice with reduced microbiota by antibiotic pre-treatment (cohort 2). Community-level and intra-species dynamics were then tracked in fecal samples over a 2-week period. As controls, samples were also collected in mice with only the colonizing *E. coli* (germ-free, cohort 3) and in mice with only the resident bacteria (cohort 4). **b**, *E. coli* bacterial load measured as colony-forming units (CFU) per gram of sampled feces for the colonized mice cohorts with innate microbiota (im), reduced microbiota (rm), and germ-free (gf). **c-e**, Frequency of the chromosomal barcodes during colonization. The most frequent 1000 barcodes are colored uniquely, whereas the rest are shown in gray. Identical barcodes are colored similarly across mouse replicates and cohorts. **f**, Effective diversity index of *E. coli* chromosomal barcodes, 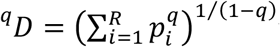 where *P_i_* is the frequency of barcode *i, R* is the total barcode count, and *q* is the order of the diversity. Effective diversity reports the count of unique barcodes (*q* = 0), frequency-weighted diversity (*q* = 1), or inverse frequency of the dominant barcode (*q* = ∞). **g,** Effective diversity for the microbiota based on the frequency of bacterial families from 16S rRNA profiling.

It is well-established that higher diversity and species richness of an ecological community makes it less susceptible^33^ to invasion including the gut microbiota. However, this resistance to invasion can be compromised upon environmental perturbations, such as antibiotic treatments, that reduce community diversity, making them susceptible even to non-pathogenic bacteria. Additionally, the gut itself presents a complex “biogeographical” environment, where distinct selective niches arise from heterogeneity in the availability of metabolites, nutrients, and immune effectors, as well as, epithelial topography and mucus architecture^34^. With these considerations in mind, we designed mice cohorts with different complexities in their gut bacterial microbiomes (Fig. 1a): mice with innate microbiome (cohort 1, “im”) and mice with reduced microbiome due to pre-treatment of antibiotics (cohort 2, “rm”); and germ-free mice (cohort 3, “gf”). Lastly, as a control for the community dynamics in the absence of colonization, we had another cohort of mice that received antibiotic treatment but was not gavaged by *E. coli* (cohort 4, “nc”). Cohorts 2 and 4 were pretreated with an antibiotic cocktail (metronidazole, neomycin, ampicillin, and vancomycin) for three weeks followed by three days of no treatment to flush out the antibiotics. On day zero, *E. coli* were introduced in mice of cohorts 1, 2, and 3. Then, for all cohorts, fecal samples were taken at 3h, 6h, 12h, and 24h on day one and then once daily for two weeks. The multiple sampling on day one was required to capture the kinetics of transit through the gut of the colonizing bacteria^29^. Extraction of bacterial genomic DNA from the feces, followed by PCR amplification of the *E. coli* barcoded region and next-generation sequencing, afforded high-resolution lineage tracking during gut colonization (Fig. 1c-e and Extended Data Fig. 1a). We also simultaneously tracked the community dynamics of resident bacteria using 16S rRNA profiling (Fig. 2b, 3b, and Extended Data Fig. 4b).

**Figure 2.**
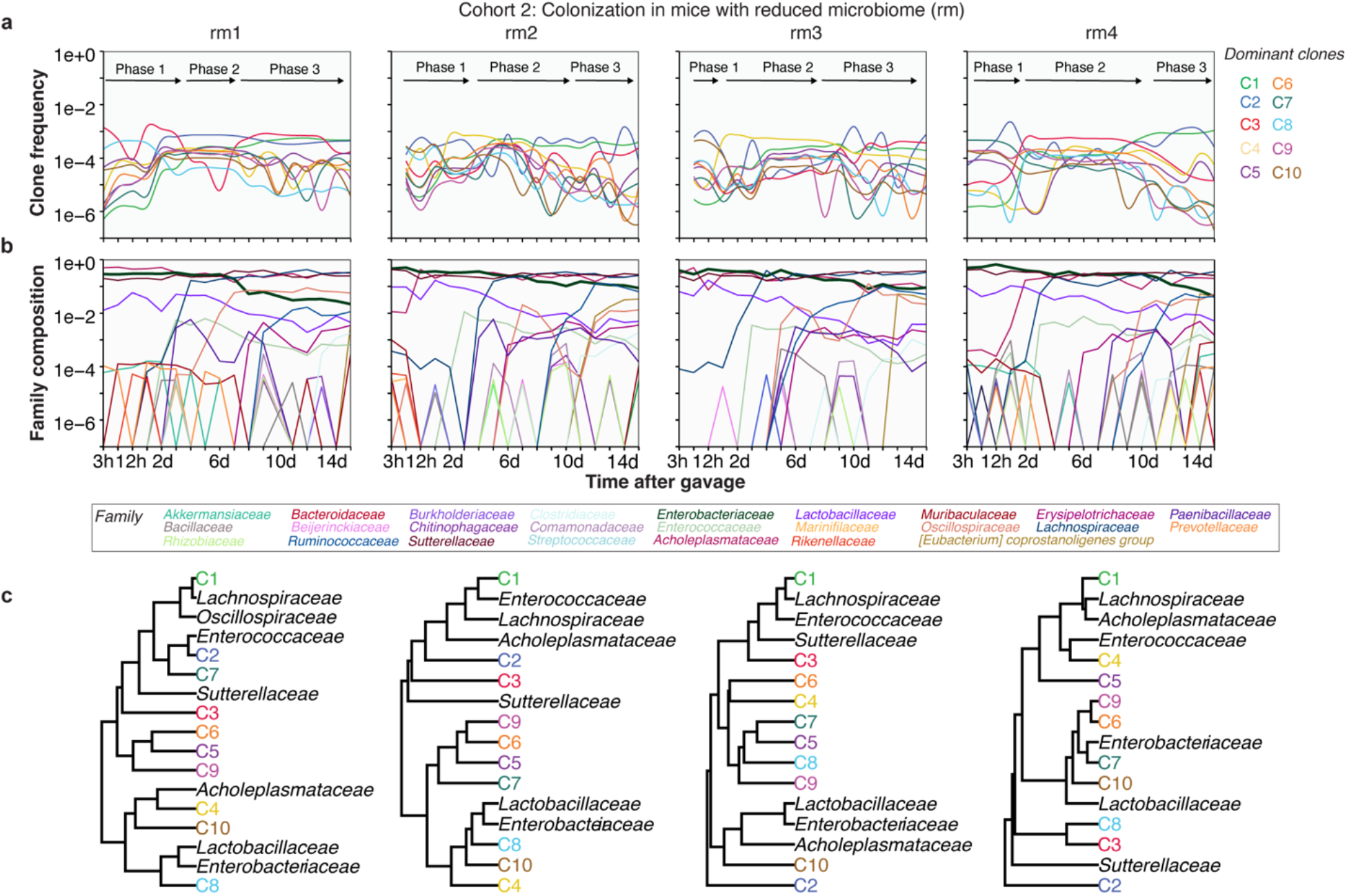
Clone-specific community interactions in the gut for the rm cohort. **a,** Dominant clones are determined by clustering of barcodes that persisted for most of the 2-week period. Pairwise pearson correlation of the barcode’s frequency time-series is used as the distance metric. These dominant barcodes represent ~5 - 7% of total unique barcodes. The putative dominant clones are ranked based on their average frequency at the end of the timeseries. **b,** Community dynamics (16S rRNA profiling) are analyzed at the level of the family. *E. coli* is a member of *Enterobacteriaceae*, which is shown as a thicker line. **c,** Co-clustering *E. coli* clones with the different bacterial families suggest clone-specific community interactions. The clone-species interactions are broadly reproducible across different mice, whereby the dynamics of C1 is related to *Lachnospiraceae*, and C8 and/or C10 are related to *Lactobacillaceae*

**Figure 3.**
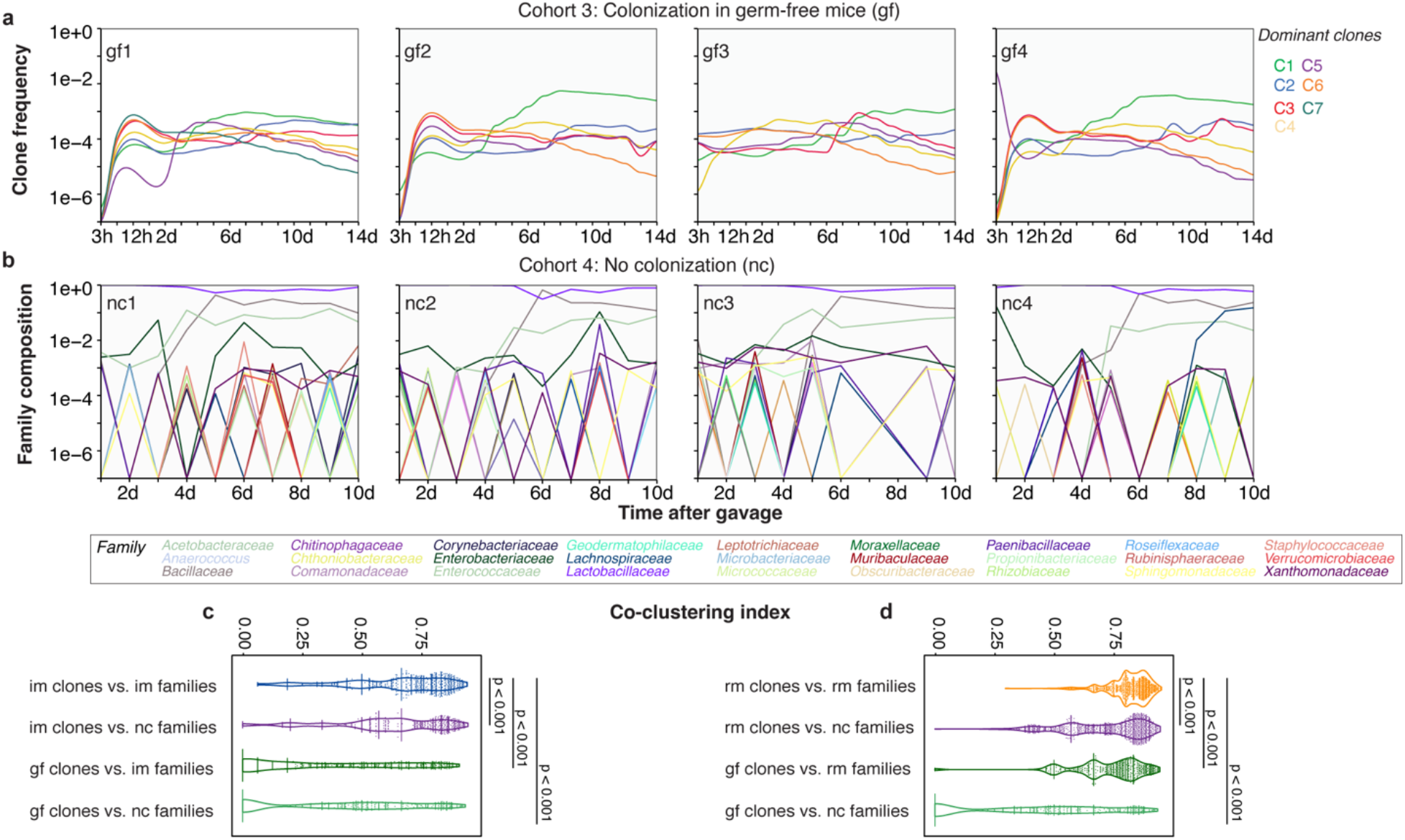
E. coli clone and bacterial community interactions are strongest when coming from the same cohort. **a,** Dominant and persistent clones of *E. coli* in pre-germ-free mice showing simpler lineage dynamics compared to cohorts 1 and 2. **b,** Community dynamics mice with reduced microbiota, but non-gavaged with E. coli, showing the recovery of bacterial community from the treatment of antibiotic cocktail. **c-d,** Co-clustering of clones and families depends on environmental conditions. Co-clustering is measured by a mixing coefficient that compares the distances in the hierarchical tree among families with distances between families and clones (Extended Data Fig. 5).

The global tempo of *E. coli* colonization and its impact on the gut microbial community is reflected by the fluctuations in the diversity of chromosomal barcodes and 16S rRNA dynamics. Thus, we calculated the effective diversity index ^*q*^*D* of the barcoded *E. coli* population that captures three complementary notions of lineage diversity. Specifically, *q* = 0 reflects the total number of unique barcodes (its “richness”), *q* = 1 is the frequency-weighted lineage diversity equal to the exponential of Shannon entropy, and *q* = ∞ reflects the reciprocal of the frequency of the most abundant barcode. The effective diversity of the chromosomal barcodes for im, rm, and gf cohort is shown in Fig. 1f. Similarly, we calculated the diversity index of the gut community composition from 16S rRNA, where the amplicon sequence variants (ASV) are defined at the level of bacterial families (Fig. 1f). Analogous to barcode diversity, *q* = 0 is the total number of unique bacterial families, *q* = 1 is the frequency-weighted family diversity, and *q* = ∞ is the reciprocal of the frequency of the most abundant family. Expectedly, the gut community of the im cohort was resistant to the invasion of the *E. coli* K12, where after an initial of ~10^6^ to ~10^8^ colony-forming units (CFU)/gram of feces within ~3h, the bacterial load reduced to below ~10^4^ CFU/gram by day 6 (Fig. 1b). The bacterial community dynamics are unperturbed by the entry of *E. coli* into the community (Fig. 1g). The inability of the *E. coli* population to establish in the im cohort is also reflected in the more rapid transit of the barcodes through the gut, whereby the peak in barcode diversity (q=0 in Fig. 1f) for mice im2-4 mice is ~3h compared to ~6h for the rm and gf mice.

In contrast, *E. coli* successfully colonized the rm cohort, reaching bacterial loads of ~10^8^ colony-forming units (CFU)/gram of feces within ~6h (Fig. 1b), which coincided with the maximal barcode diversity (Fig. 1f). The absence of resident bacteria in the gf cohort resulted in *E. coli* reaching a higher bacterial load of ~10^10^ CFU/gram of sample within ~6h. Interestingly, despite the difference in their CFU levels (Fig. 1b), both the gf and rm cohorts reach optimal diversity in ~6h (Fig. 1e). Moreover, even the unsuccessful colonization of the im cohort reached an optimal diversity around 3h, and the diversity is overlapping with the im, rm, and gf cohort. Altogether, these results suggest that the very early dynamics of entry of *E. coli* (~6 h) is primarily dictated by adaptation to the biogeography of the mouse gut rather than by the diversity of the microbial community. After this time-point, the different cohorts exhibited distinct behavior for their diversity (Fig. 1f,g).

Notably, the CFU counts for mice in the same cohort are broadly indistinguishable (Fig. 1b) despite the complexity of the underlying clonal dynamics viewed at higher resolution (Fig. 1c-e). Additionally, although barcodes begin appearing within ~3h (Fig. 1e), they were not observed in the CFU counts for the rm and gf cohorts (Fig. 1b). Barcodes that appeared first were not always the dominant ones at the end (Extended Data Fig. 1a). This observation is most drastic in the im cohort. Three mice (im1, im2, and im3) showed a reduction in barcode frequency accompanying the drop in CFU but manifested an increase in diversity towards day 6 (Fig. 1b,f). One mouse (im4) maintained a high barcode diversity despite the drop in CFU. These results highlight the stochasticity of transmission kinetics through the intestinal gut’s distinct “island” niches^34^, which is not reflected simply by measuring the total bacterial count of the invading species.

In accordance with previous lower-resolution colonization experiments^29,35^, we observed soft clonal sweeps of barcodes that eventually became dominant (>5%) after ~2 days (Extended Data Fig. 1b, rm mice; Extended Data Fig. 1c, gf mice). This clonal sweep was stronger in the gf than in the rm mice, as only one barcode reached >5% in the rm, whereas several dominant barcodes co-segregated over two weeks in the gf. These dominant barcodes originated at frequencies around 10^-7^ (Fig. 1d-e). The sweep in the gf and rm mice is manifested by a drop in barcode frequency-dependent diversity (q=1 in Fig. 1f). The drop in diversity occurs earlier in the rm cohort (~2 days), compared to gf (~7 days in *q* = 1, ~4 days in *q* = ∞). Interestingly, despite the earlier clonal diversity drop in rm, which coincides with the resurgence in bacterial community diversity (Fig. 1g), rm maintains a greater clonal diversity compared to the gf towards the end of the 2-week period (Fig. 1f). This suggests that gf, adjusting only to the selective pressure imposed by gut biogeography, is subject to a stronger intra-species competition. In contrast, the resurgence of the resident bacterial community in the case of rm (Fig. 1g) leads to the coexistence and co-dominance of multiple barcodes in the *E. coli* population (Fig. 1f). Altogether, this highlights the interaction of the resident community and the clonal lineages of the invading species.

### Clone-specific interaction between *E. coli* and other bacteria

Next, we determined the impact of the colonizing species on the composition of the resident bacterial community. While the community in the im cohort was more resistant to invasion, the introduction of *E. coli* in the gut microbiome led to a reduction in the abundance of some resident bacterial communities in the rm cohort (16S rRNA in Fig. 2b and 16S rRNA in Extended Data Fig. 4b). This quick initial collapse happens within the first ~3h and is clearly manifested in rm2 (16S rRNA in Fig. 2b). However, the establishment of *E. coli* was accompanied by the resurgence of the bacterial community around day 4, followed by the coexistence of *E. coli* and the resident bacterial community. Notably, *Sutterellaceae* was unperturbed by the introduction of *E. coli* in all four mice. Although *Muribaculaceae* was unperturbed only in rm 1 and rm 3, it was the first bacterial family to rebound in rm 2 and rm 3 (Fig. 2b). The canonical member of a gut microbiome, *Lactobacillaceae*, had an intermediate abundance when *E. coli* was introduced, but this declined after *E. coli* established in the gut microbiome. In the no-colonization cohort (nc), the community also exhibited a rebound due to the release from the antibiotic treatment (Fig. 1g and 3b). However, the resident community of the nc cohort was dominated by *Lactobacillaceae* and was less diverse than the resident community of the rm cohort. These results demonstrated the impact of *E. coli* introduction on bacterial community composition.

To determine if there are interactions between specific *E. coli* clones and bacterial families, we first identified the dominant *E. coli* lineages using barcode dynamics (Fig. 1c-e). The barcode lineage dynamics reflects its effective fitness (selection coefficient) over time^32^, and thus similarities in barcode lineage dynamics can be indicative of *E. coli* clones under similar selection coefficients. We performed a shape-based clustering analysis of linage dynamics using the Pearson correlation as the similarity measure computed for pairs of barcodes with mean frequency >5e^-5^ and persisted for at least 12 of the 18-time points (Extended Data Fig. 3). These represent ~5-10% of the total barcoded *E. coli* observed in the rm and gf mice (Extended Data Fig. 3b,c). Clusters of persistent barcodes were defined as putative clonal lineages hereafter and ranked based on average frequency (Extended Data Fig. 3). Interestingly, in this procedure, the clonal cluster C1 always contained the dominant barcode lineages that exhibited the sweeps, even if C1 cluster itself did not have the largest number of barcodes (Extended Data Fig. 3). This observation validates the lineage clustering approach. A LOESS regression of the top 10 clonal clusters of *E. coli* highlights the persistence of extremely rare barcodes in the colonizing population (Fig. 2a & Fig. 3a)

Considering the diversity of these clonal lineage clusters, we asked whether individual clusters were associated and potentially interacting with specific bacterial families. We performed co-clustering of the dynamics of putative clonal lineages of *E. coli* and those of the 16S rRNA bacterial community profiles using the k-shaped algorithm^36^ (Fig. 2c, Extended Data Fig. 4c Extended Data Fig. 5). These analyses revealed a consistent picture across the rm cohort. The dominant cluster, C1, was always grouped with *Lachnospiraceae*, whereas two other low-frequency clusters, C7 and C8 grouped persistently with *Lactobacillaceae*, the canonical member of gut microbiota (Fig. 2c). Interestingly, it was previously shown in invasion studies of pathogenic strains of *E. coli* and *Lachnospiraceae* that these bacteria utilize similar sugars and thrive in the same environment^37^.

To demonstrate that the degree of co-clustering of *E. coli* clonal lineages and bacterial families was specific to the rm cohort, we applied the shaped-based co-clustering of barcode lineages for the gf (Fig. 3a) and the nc cohort (Fig. 3b). The extent of co-clustering was measured using the mixing index, *D_c,m_* = 1 — (max|*F*(*c*) – *F*(*m*)|) (Fig. 3c-d and Extended Data Fig. 5), which compares the clustering distance from clonal lineages to bacterial families *F(m*) with the distance between clonal lineages *F*(*c*) (Fig. 3d). Indeed, the co-clustering between clones and bacterial families was strongest in the rm cohort(Fig. 3d). Expectedly, co-clustering was weakest when the 16S rRNA community dynamics of im and rm were paired with the gf cohort clonal lineages (Fig. 3d).

Moreover, we applied the same analyses to the im cohort to test if there is a similar heterogeneous interaction between the invading *E. coli* population and the bacterial community, even if the invasion was unsuccessful. Indeed, the shaped-based co-clustering showed heterogenous interaction between *E. coli* lineages and bacterial families (Extended Data Fig. 4). The extent of co-clustering in the im cohort is weaker than in rm (Fig. 3c-d, im is blue, rm is orange), which is in agreement with the resilience of the im community to the invasion. However, despite the unsuccessful E. coli invasion in the im mice, the co-clustering between E. coli clonal lineages and bacterial families is strongest when they come from the same biological cohort (Fig. 3c), suggesting intra- and inter-species interactions (Extended Data Fig. 4c), even if only transient. Altogether, the high-resolution lineage tracking demonstrated that intraspecific variation could lead to clone-specific interactions in bacterial communities during invasions, including those that may not lead to establishment.

### Dynamic covariance mapping defines phases of colonization

The barcode and 16S rRNA time series suggest distinct phases in the colonization dynamics in rm mice (Fig. 2a-b). Specifically, the entry of the invading *E. coli* population is manifested by the rise of the dominant clone frequency in phase 1. This is accompanied by an increase in CFU and barcode diversity (Fig. 1b,f) and a drop in resident bacterial community diversity (Fig. 1g and Fig. 2b). In phase 2, relative stasis and lower fluctuations in the dominant clone frequencies as the resident bacterial community re-emerges. Lastly, in phase 3, the dominant clones of *E. coli* coexisting with the resident community undergo large fluctuations. These phases are distinct from the much simpler dynamics of clonal lineages observed in the gf cohort (Fig. 3a), which suggests that the phases in rm are driven by interactions between the colonizing *E. coli* and the resident community.

To unambiguously define these phases and the dynamic stability of the community, we developed the method of dynamic covariance mapping (DCM) that estimates the time-dependent community matrix of pairwise interactions at the level clones. Specifically, the community dynamics can be described by a vector of time-series, *Z*(*t*) = {*z*_1_(*t*),*z*_2_(*t*),...}^*T*^, featuring 10 time-series corresponding to the dominant *E. coli* clones and 7 for bacterial families. Although the full time evolution of *z*(*t*) is complex and unknown ^2,9,38,39^, the Jacobian matrix *J* of its linearized dynamical system, representing the community matrix of pairwise interactions, can be estimated as the covariance of one time-series and another’s time derivative^9^, *J* = 〈cov(*ż_i_,z_j_*)〉. The eigenvalues of the Jacobian matrix report on the dynamics and stability of the system^2,38,39^. When the real part of an eigenvalue is negative, the system is stable against perturbation along the direction of the associated eigenvector, and conversely, when positive, the system becomes unstable. The magnitude of the imaginary part of an eigenvalue implies oscillatory behaviors.

We quantified the Jacobian over a progressively shifting time window and performed eigenvalue decomposition (Fig. 4, Supplementary Movie 1a-d). The real parts of eigenvalues corresponding to early time-points were positive, reflecting that the system is unstable due to the introduction of *E. coli*. This is accompanied by the drop in bacterial community diversity (Fig. 4c left panel). This was followed by a second phase, with the real parts of eigenvalues moving from positive to negative, corresponding to the recovery of system stability and the resurgence of some bacterial families (Fig. 4a, c (middle panel)). Finally, in phase 3, the system became dynamically stable but with notable oscillations in both the 16S and the clonal dynamics (Fig. 4c, (right panel)). These results highlight the rapid time dependence of the community interaction matrix.

**Figure 4.**
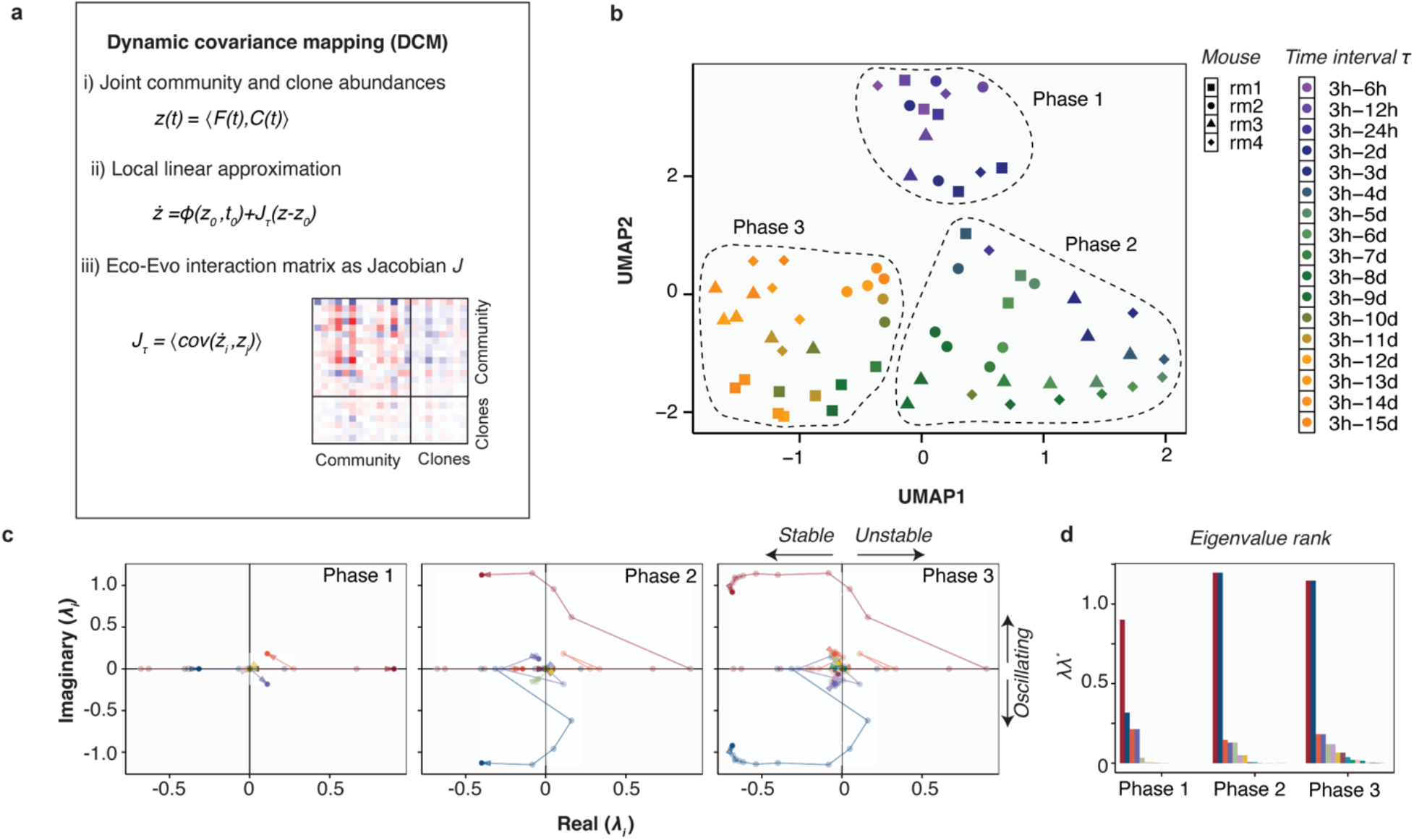
Distinct phases of colonization quantified by the Dynamic Covariance Mapping. **a,** Dynamic covariance mapping (DCM). i) The time series *z*(*t*) is a concatenated vector of the family and *E. coli* clone frequencies. ii) The evolution of *z*(*t*) is governed by a system of ODEs that can be linearly approximated by the Jacobian matrix *J_τ_*, which is the co-variance between the time-series *i* and the time-derivative of the time-series *j*. The Jacobian is calculated over a time interval *τ. t*_0_ is the start of the experiment. iii) The expanded interaction matrix includes interfamily, intra-clone, and family-clone interactions. **b,** 2D UMAP projection of the eigenvalues of the time-dependent covariance matrix *J_τ_*. The eigenvalues cluster into three distinct time domains that reflect the phases of colonization shown in Fig. 2a. **c,** Stability analysis over the three phases in Fig. 2a mouse rm1. Phase I is transient instability corresponding to the entry of *E. coli* and the collapse of resident bacteria. Phase II is the return to a stable regime and the re-emergence of the community species. Phase III is quasi-dynamic equilibrium with both oscillations in the clonal and community dynamics (see Supplementary Movie1a-d). **d,** Time-varying eigenvalues of the mouse rm1 were ranked and colored according to their magnitude. Ranking revealed that the first four eigenvalues dominate the whole dynamics.

We then determined how reproducible and consistent these three phases are across the mice cohorts (Supplementary Movie 1a-d). For each time interval, there are ~17 eigenvalues corresponding to the Jacobian matrix. Using a Uniform Manifold Approximation and Projection (UMAP) (Fig. 4b), we found that the eigenvalues as a function of time broadly overlap across different mice, with the boundaries of the 3 phases unambiguously determined (unbiased clustering in the UMAP; Extended Data Fig. 6d). Interestingly, application of DCM to the unsuccessful invasion in the im mice also revealed two distinct phases for im1-3 (Extended Data Fig. 6g), corresponding to the entry of barcoded E. coli (~3h to 12h) followed by the collapse in barcode diversity due dramatic drop in bacterial load (Fig. 1b,f and Extended Data Fig. 1a).

Are these distinct temporal phases primarily driven by the 16S, clonal dynamics, or both? To answer this question, we combined all eigenvalues from the clone dynamics of the rm and gf cohort and projected them onto a single UMAP (Extended Data Fig. 6h). This revealed that the entire 2-week dynamics of *E. coli* in the gf cohort and rm cohort are not overlapping at all. Also, rm clone DCM analysis revealed that without the bacterial community dynamics effect, they cluster within the mouse rather than the cohort. This suggests that the *E. coli* clonal dynamics in the rm cohort were largely driven by the interaction between clones and the bacterial community.

### Estimates of the relative fitness of the gut community during the 3 phases of colonization

Since the gut is a multi-species system, the effective fitness manifested by clones or bacterial families reflects their adaptation to the mouse’s intestinal biogeography, interactions with other species and clones, and impacts from mutations. This complexity reflects the fundamental coupling of evolutionary and ecological forces. To determine how the 3 phases defined from DCM correspond to the fitness experienced by *E. coli’s* clones and bacterial families, we estimated the relative per-capita growth rate of the clones from the time derivative of their normalized frequency (Methods). Within the rm cohort, we partitioned these relative fitness estimates according to the three DCM-identified phases and found that the dominant clone cluster C1 experienced a positive fitness as the *E. coli* population adapts to the gut biogeography (>50% of relative fitness is positive; Fig. 5a). Similarly, its most dominant interactor (Fig. 2b), *Lachnospiraceae*, was also driven by positive fitness(Fig. 5d). Experienced relative fitness was symmetric for the next two dominant clones C2 and C3. The relative fitness experienced by low segregating clones shifted from positive in phase 1 to negative in the other phases (Fig. 5b).

**Figure 5.**
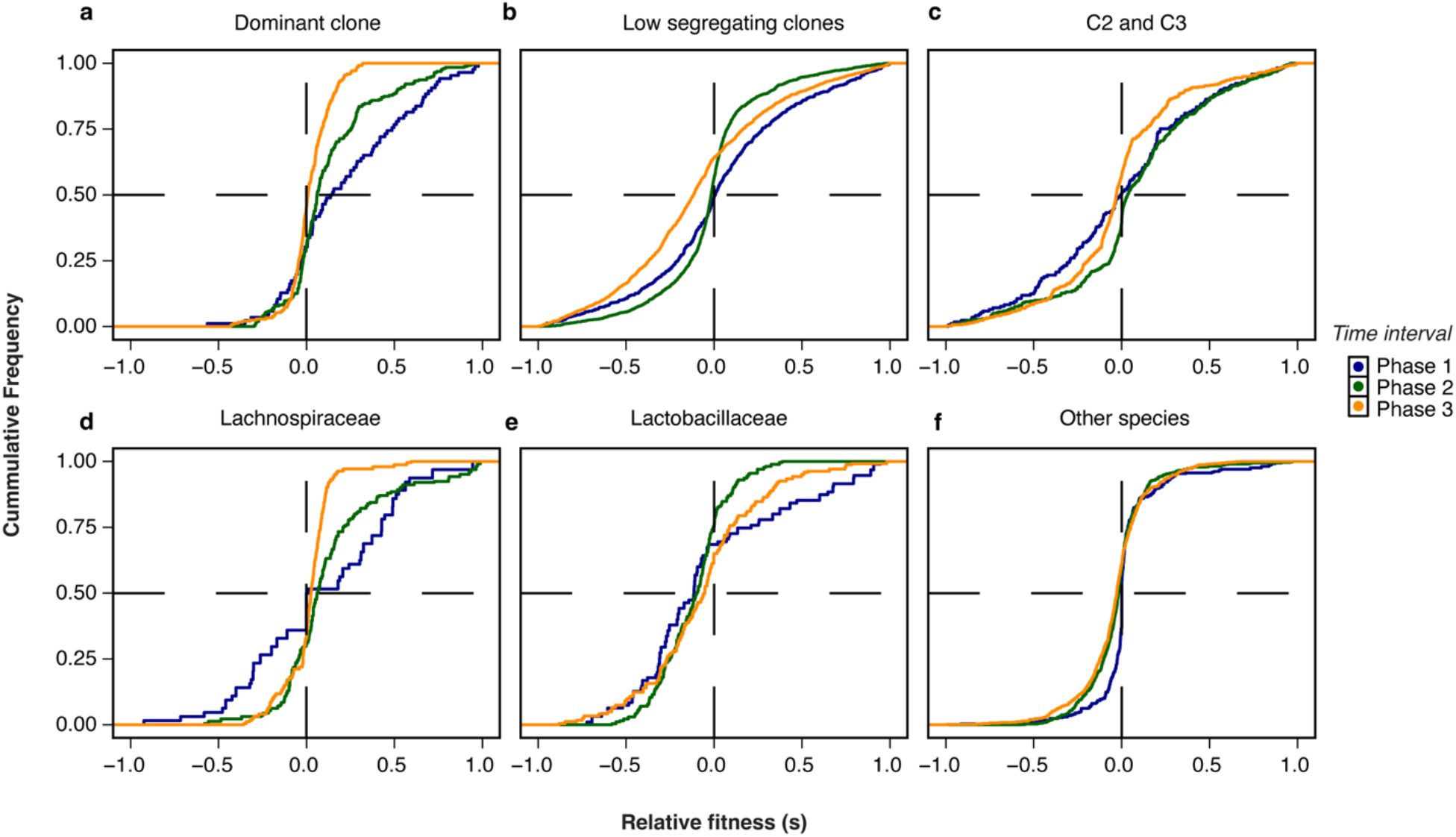
Relative fitness of the dominant clones and bacterial families. **a-f,** The relative fitness is estimated from the slope of the clone and family frequency. The cumulative distribution is defined over the 3 phases in Fig. 4. Vertical dashed lines indicate neutral (s=0). Horizontal dashed lines indicate 50%. The dominant clone C1 is primarily driven by adaptive changes in phase 1 and then reverts to equal fractions of deleterious and beneficial mutations in phase 3, which is consistent with dynamic equilibrium. Fitness at the community level shows that the dominant interactor of C1, *Lachnospiraceae*, is also experiencing strong adaptive changes in phase 1 and equal fractions of beneficial and deleterious fitness effects in phase 3.

The system reached a quasi-steady state where there was an equal fraction of positive and negative fitness changes (Fig. 5c orange curves for dominant clones C1, C2, and C3), corroborating the eigenvalue decomposition analyses, which indicates a stable oscillator in phase 3. In single-species systems, this implies a mutation-selection balance whereby there is an equal fraction of beneficial and deleterious mutations. However, in species-rich communities, the oscillation is primarily driven by co-evolution from clone-specific interactions. Indeed, the estimated fitness for the gf cohort (Extended Data Fig. 6) revealed minimal fitness changes, suggesting that the contribution of *de novo* mutations in the 2-week period was less compared to the rm cohort with complex interactions in resident bacterial communities.

### The dynamic similarity in phase 1 is driven by similar barcodes

How similar are the clones across different mice, and are they driven by the same barcodes? To this end, we performed pairwise clustering using the Pearson correlation as the similarity of the clone time-series. In both rm and gf cohorts, we observed that the dominant clonal lineages (C1 and, to a lesser extent, C2) have similar time series (Fig. 6a, d). By calculating the overlap coefficient between barcodes in each clone, we found that the dominant clonal lineages are more likely to be the same barcodes (Fig. 6e). Considering that the dominant clones in the rm cohort change their dynamics primarily in phase 1 (Fig. 4), this suggests that the reproducibility of dominant barcode dynamics and their consistent interaction with *Lachnospiraceae* is likely driven by standing genetic variation in the colonizing population. That is, the gavage *E. coli* pool has genetic variation^24^ which drives its early adaptation dynamics in the gut. In further support of this proposition, the effect of standing genetic variation was strongest in the colonization of gf mice (Fig. 6b), where the similarity in clonal dynamics across mice was driven by strong barcode similarity.

**Figure 6.**
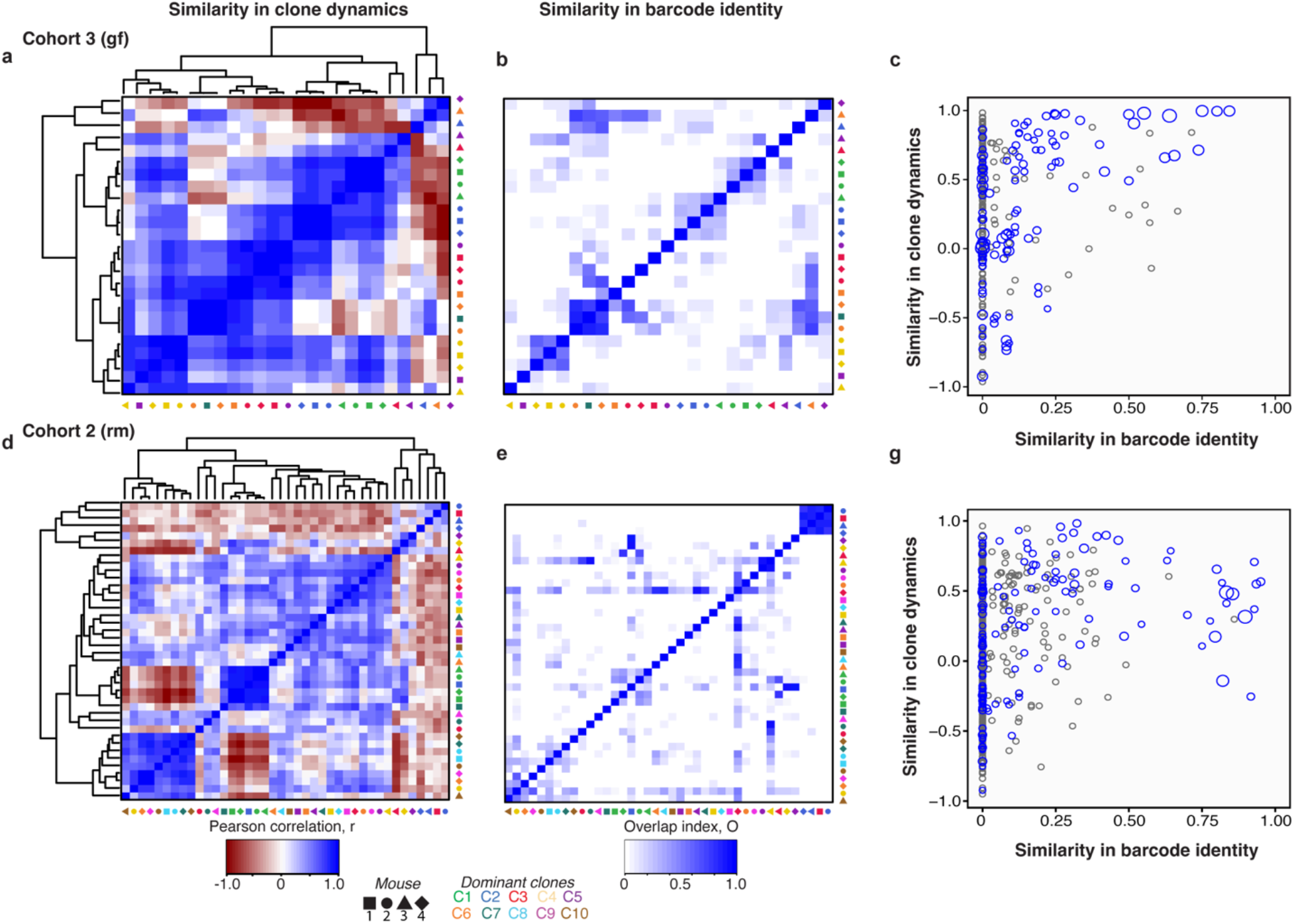
The similarity of clonal dynamics across mice is partly driven by identical barcodes. **a**, Similarity between the time series of dominant clones across all 4 gf mice quantified by Pearson correlation. Matrix elements are clustered based on hierarchy (dendrograms indicated). Colors indicate the clone’s identity, while the shape indicates the mouse of origin. **b,** The similarity in barcode identity between the different clones is quantified by the overlap coefficient, *OC*(*A, B*) = |*A* ⋂ *B*|/*min*(|*A*|, |*B*|), where A and B are the sets of unique raw DNA barcodes that belong to two dominant clones. Identities of matrix elements are similar to panel **a**. **c,** Scatter plot of the similarity in dynamics between two clones by Pearson correlation vs. similarity in their barcode identity by overlap coefficient. Overlap coefficients satisfying p-value <0.05 from bootstrapping are shown in blue; otherwise, they are shown in grey (Methods). The size of the circle is proportional to the significance of the overlap coefficient. **d-g,** Similarity in dynamics and barcode identity for the colonization in mice with the resident microbiome.

In the rm cohort, for clonal lineages that were less dominant but exhibited persistent oscillations in phase 3 (Fig. 5), their similarity in barcode dynamics was not accompanied by similarity in barcode identity. This indicates that even within 2 weeks, the clonal lineages in a mouse gut start to diversify in phase 3, and according to relative fitness, this diversification in dynamics is more likely driven by ecological effects (e.g., stochasticity in the interaction with resident bacterial species) than *de novo* mutations.

## Discussion

Our experimental and computational framework offers a generalized approach to quantify microbial community interaction matrix and its consequences on dynamic and stability, particularly following perturbations triggered by invading species. With our experimental barcoding protocol, we demonstrate that intra-species variation leads to time-dependent interactions, even during the early stages of community colonization. Although the dynamics are complex, the global colonization dynamics are surprisingly replicable and can be defined by 3 phases that arise from the coupling of ecological and evolutionary dynamics. This seems contradictory to other studies showing a lack of reproducibility and replicability of microbiome composition across mice replicates^40^. However, we note the onset of divergence between the mice cohort (Fig. 4b) after 2 weeks due to stochasticity of *de novo* mutations. We cannot yet comment on the long-term implication of ISV at the resolution afforded by this experiment since our barcode diversity is exhausted after a clonal sweep. This would require a “renewal” or regeneration of new DNA barcodes, as recently done in yeast^41^.

Additionally, intra-species diversity is present not only in the colonizing species but also in the resident community; thus, our chromosomal barcoding approach could be extended to species that are innate to the gut microbiota. The high-resolution colonization dynamics could also be extended by pathogenic barcoding species, such as *P. aerogenousa* and *S. enterica*, which are more aggressive colonizers than *E. coli* K12. Therefore, we argue that the gut microbiome for us is an ecological system, such that all the approaches presented here could be broadly applicable to most microbial ecological networks. However, the gut microbiome has particularities. More specifically, the gut microbiota itself is shaped by the genetics and phenotypes of the mice, which we do not explore in this study. Indeed, the mice themselves, in general, are not homogenous and could have an impact on the gut composition. In human microbiomes, it was shown that genetic variation in humans could itself impact the diversity of the microbiomes^42^. In the future, the impact of host diversity will be explored by performing colonization experiments on mice with diverse genetic backgrounds.

Broadly, the DCM that we developed here represents a model- and parameter-free approach to analyzing the stability and distinct temporal phases of a microbial system, starting simply from high-resolution time series abundance data. Our result showed that these phases of invasion and the intra- and inter-species interactions are highly reproducible among mice replicates is rather unexpected considering the variability in microbiome compositions which is the norm in the microbiome field^43^. We argue that although specific compositions may be highly variable across mice, the overall tempo of ecological and evolutionary dynamics, as manifested by the DCM analysis, are more reproducible features of the microbiota. To this end, the DCM and its future incarnations could provide a framework for predicting the microbiota’s response to perturbations, especially in the context of the invasion of pathogenic species^44^ and fecal transplant to treat human disorders^45^.

## Methods

### Experimental procedures

#### (i) *E. coli* barcoded population generation

Barcoded *E. coli* populations were generated as previously described^24^ using the Tn7 transposon library. The first step is transforming the recipient *E. coli* MG1655 cells with the Tn7 helper plasmid and induction of the transposase integration machinery. The second step is the transformation of the Tn7 integration plasmid library, which integrates the barcodes into the chromosome of the bacteria. The Tn7 integration plasmids with barcode and spectinomycin cassette were extracted from TransforMax EC100D pir + cells (Lucigen) with a Qiagen midi kit. Then *E. coli* MG165 cells were transformed with the Tn7 helper plasmid to induce the transposase integration machinery. Transformed cells with Tn7 helper plasmid were grown overnight in LB supplemented with 100μg/ml ampicillin at 30 °C. In these cells, transposon machinery was induced with arabinose to transform with Tn7 integration plasmids. After overnight incubation on the bench, they were plated on LB agar plates containing 100 μg/ml spectinomycin. Randomly picked colonies were checked for chromosomal incorporation of barcode cassettes by targeting the Tn7 integration site. We scraped all the colonies from the plates, then pooled, thoroughly mixed, and aliquoted them with 15% glycerol. These stocks were stored at −80 °C pending the mice colonization experiments.

#### (ii) Mice evolution experiments

We used several cohorts of mice to determine colonization dynamics in their gut: Cohort 1 (im) mice with <u>i</u>nnate <u>m</u>icrobiota followed by *E. coli* colonization (4 replicates); Cohort 2 (rm) or mice with reduced <u>m</u>icrobiota and pre-treated with an antibiotic cocktail followed by *E. coli* colonization (4 replicates); Cohort 3 (gf) or mice that were initially <u>g</u>erm-<u>f</u>ree and colonized with barcoded *E. coli* barcode (4 replicates); and Cohort 4 (nc) or mice with microbiota and pre-treated with an antibiotic cocktail but <u>n</u>ot <u>c</u>olonized by *E. coli* (4 replicates). Cohorts 2 and 4 (rm and nc) were administered an antibiotic cocktail (metronidazole 1 g/L, neomycin 1g/L, ampicillin 1g/L, and vancomycin 0.5 g/L) for four weeks to reduce the complexity of the gut microbiota. Under these conditions, 99.5% of the cecal bacteria are eliminated at the end of treatment^46,47^. Then, we let them recover for three days *without antibiotics* before introducing the barcoded population, which we set as our day zero. After gavage of the barcoded population, fecal samples were taken at 3, 6, 12, and 24 hours and once daily until day 14 for rm and day 15 for gf. The nc cohort fecal samples were collected for ten days. During the day of fecal collection, we split the sample, one for bacterial load measurements (see below) and another for storage at −80 °C until subsequent genomic analysis. 80 μl of the feces homogenate was placed with 20 μl of 100% glycerol to make 20% glycerol stocks for later recovery of live bacteria.

#### (iii) Bacterial load measurement

To measure the bacterial load in the fecal samples, we spread them with increasing dilutions on LB plates with spectinomycin 50 μg/ml to select for the colonizing *E. coli*. The chromosomal barcode contains the spectinomycin resistance cassette (spR)^24^. Measurements of bacterial loads were done in 3 independent replicates.

#### (iv) Genomic DNA extraction in fecal samples, chromosomal barcode amplification, and next-generation sequencing

Genomic DNA (gDNA) was extracted from whole fecal pellets using the QIAamp Fast DNA Stool Mini kit (Cat: 51604). A two-step PCR was used to amplify the chromosomal barcodes and then append the Illumina adapter sequences. For the first PCR, anywhere between 20 to 100 ng of template per sample was used with PrimeSTAR GXL DNA Polymerase from TAKARA (Cat: R050B). The parameters for this 1^st^ reaction were as follows: 94 °C for 5 min, 30X (95 °C for 10s, 53 °C for 15 s, 68 °C for 45 s), 68 °C for 5 min, hold at 4 °C. The Primers for this PCR are the following: 5’-TCGTCGGCAGCGTCAGATGTGTATAAGAGACAG-3’, 5’-GTCTCGTGG GCTCGGAGATGTGTATAAGAGACAG-3’. The resulting amplicon sequence from this PCR is the following: 5’ -gatatcggatcctagtaagccacgttttaattaatcagatccctcaatagccacaacaactggcgggcaaacagtc gttgctgattggtcgtcggcagcgtcagatgtgtataagagacagtcgcgccggNNNNNNNNNNNNNNNtatctcggtagtgggatacgacgataccgaagacagctcatgttatatcccgccgttaaccaccatcaaacaggattttcgcctgctggggcaaaccagcgtggaccgcttgctgcaactctctcagggccaggcggtgaagggcaatcagctgttgcccgtctcactggtgaaaagaaaaaccaccctggcgcccaatacgcaaaccgcctctccccgcgcgttggccgattcattaatgcagctggcacgacaggtttcccctgtctcttatacacatctccgagcccacgagacgccactcgagttatttgccgactaccttggtgatctcgcctttcacgtag-3’. The contiguous 15 *N*s in this amplicon sequence corresponds to the random nucleotides that serve as our chromosomal barcodes^24^. The product from this PCR was purified and cleaned with NucleoSpin Gel and PCR clean-up kit from TAKARA. A 2^nd^ PCR was performed with high-fidelity PrimeSTAR GXL DNA Polymerase (Takara Cat: R050B) to add the Nextera indices (Nextera XT primers Set A 96 Indexes, 384 Samples, Cat# FC-131-2001). We followed the suggested cycling conditions, which are as follows: 94 °C for 5 min, 12X (95 °C for 10 s, 55 °C for 15 s, 68 °C for 45 s), 68 °C for 5 min, hold at 4 °C. The primers for this 2^nd^ reaction were the following: 5’CAAGCAGAAGACGGCATACGAGAT[I7]GTCTCGTGGGCTCGG-3’ and 5’-AATGATACGGCGACCACCGAGATCTACAC[I5]TCGTCGGCAGCGTC-3’. PCR products from all reaction tubes were purified with magnetic beads (Beckman Coulter) and pooled together, spiked with 15% of PhiX DNA, and sequenced using either Miseq or Nextseq Illlumina chips at Université of Montréal’s IRIC Genomic Platform. Bioinformatic analyses are described in the Analysis section below.

#### (v) 16S profiling

Similar to the chromosomal barcode amplification, we used a two-step PCR to amplify the genomic region of interest and prepare the library for Illumina sequencing. The 16S rRNA V4 region was PCR-amplified with buffer and polymerase PrimeSTAR GXL DNA Polymerase (Takara, Cat: R050B). The cycling conditions for the PCR are as follows:_98 °C for 3 min, 35X (95 °C for 10 s, 60 °C for 15 s, 68 °C for 35 s), 68 °C for 5 min, hold at 4 °C. The primers for the reaction are the following: <u>5_-</u>TCGTCGGCAGCGTCAGATGTGTATAAGAGACAGYRYRGT GCCAGCMGCCGCGGTAA-3’ and 5’-GTCTCGTGGGCTCGGAGATGTGTATAAGAGACA GGGACTACHVGGGTWTCTAAT-3’. PCR products were purified with Nucleospin Gel and a PCR purification kit from TAKARA (Cat: 740609). Illumina sequencing adaptors were added to respective samples with PCR using the same primers and protocols similar to the barcode amplification. The PCR amplicons of the samples were then pooled after a purification and concentration equalization process with the AMPureXP Kit (Beckman Coulter). The libraries were processed in an Illumina MiSeq v2 (500 cycles and paired-end).

### Analysis

#### (i) Barcode extraction from the FASTQ file and determining putative “true” lineages

To understand how clonal populations of cells change over time, we first identified and extracted the barcode sequences from our raw sequencing data. To prepare reads for extraction, we prepended them with one N and a corresponding ‘?’ quality score. This was required to extract barcodes from the reads using the *bartender_extractor_com* component from the tool *Bartender*^48^. We discarded reads with an average *Phred* quality score below 30 (corresponding to the ‘?’ character) and kept reads with at most one mismatch in the sequence following the variable region, which is [TATC]. For each remaining read, a raw 15-nucleotide sequence barcode was extracted. However, not all of these raw barcodes match the true synthesized barcodes due to mutations in the sequencing and/or PCR. To correct for sequencing errors in the raw barcodes, we used the *bartender_single_com* on the raw barcodes with default settings. Here, it was assumed that an infrequent barcode with one or two mismatches from a frequent barcode was a mutant of the more frequent barcode and hence, added to the latter. This step produced a list of putative barcode lineages for the sample. Additionally, since we have multiple time points per mouse, we wanted to ensure that barcode identities were consistent across the biological samples. Thus, we pooled all the raw barcodes from the same mouse as a single list. We then applied the same *bartender_single_com* procedure to the pooled list. This step resulted in a comprehensive list of raw barcode sequences mapped to their consensus sequence for all samples from one mouse. From this list, we iteratively mapped each raw barcode sequence against all individual samples to yield the number of reads per time point per barcode lineage. For each mouse, we sequentially assigned a numeric ID to the barcode lineage to produce a list of barcode lineage trajectories for one mouse. This analysis pipeline is available on github (https://github.com/melisgncl/high-resolution-mouse-barcoding.git).

#### (ii) Visualizing barcode dynamics

To compare barcode trajectories within and between mice cohorts, we aimed to use consistent color coding for barcode lineages. First, we assigned a unique color to all lineages that reached a relative frequency of 5e-05 in their respective mouse. The frequency *f_i_*(*k*) of barcode lineage *k* in condition *i* is:

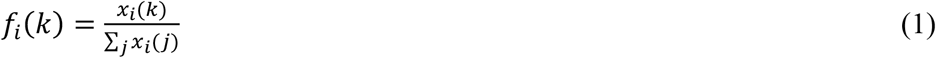

where *x_i_* (*k*) is the barcode read count. This operation was applied to each mouse, such that the color scheme was consistent when the dynamics were compared (Fig. 1c-e and Extended Data Fig. 1). For example, a barcode lineage that was assigned the color “magenta [#c20078]” will always have this color in all the figures. Conversely, no other barcode was assigned the same color. To create the Muller-type plots for each mouse (Extended Data Fig. 1), the barcode frequencies at every time point were represented in linear scale. In each mouse, the barcodes were sorted by the maximum frequency they attained over the time-series. This produced a stacked area plot where dominant barcodes were shown starting from the bottom of the panel and progressively lower-frequency barcodes were shown at the top. The same data was used to plot the frequency trajectories in log10-transformation (Fig. 1c-e). Barcodes that reached a minimum frequency of 1e-05 throughout its time-series were shown in color, whereas the remaining barcodes were shown in grey for clarity.

#### (iii) Quantification of barcode diversity

The simplest way to quantify the diversity of barcoded lineages in a population is to count the number of unique barcodes observed at a particular time point (Fig. 1c-e). However, if lineages differ widely in frequency, then this measure may not be very informative and will suffer from substantial sampling bias (since very low-frequency barcodes will be under-sampled). A more general approach is to quantify the diversity of barcodes using the effective diversity index^49^

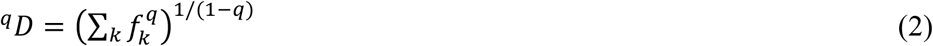

where *f_k_* is the frequency of the *k*th barcode lineage, and *q* is the “order” of the diversity index. When *q* = 0, the index simply counts the absolute diversity in the sample, i.e., the total number of unique barcode lineage. This measure is equivalent to the species richness used in ecological studies ^50^. When *q* = 1, the index weights each barcode lineage by its frequency. This measure is equivalent to the exponential of the Shannon entropy *H*;

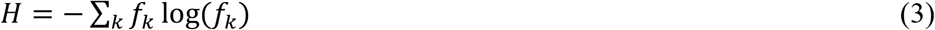

When *q* → ∞, the index is equal to the reciprocal of the proportional abundance of the most common barcode lineages. Thus, only the higher-frequency lineages contribute to the value of this index. By comparing the diversity index across these three orders for *q*, we could describe the complex dynamics of the barcode composition over the course of the experiment. In the trivial case when all barcode frequencies were equal, the effective diversity index would be equal to the absolute number of barcodes regardless of the order of *q*. We should expect absolute diversity (*q* = 0) to be no greater than the maximum theoretical diversity of the barcode library. Additionally, we should also expect this measure to decrease over time as barcodes are lost from the population since diversity is exhausted In time within host transit and dynamics.

#### (iv) Barcode lineage clustering

To identify the clonal lineages, we clustered the barcode lineages for each mouse based on the similarity of their time series behavior. To maximize the accuracy of this clustering, we excluded barcodes with insufficient time points. Specifically, for each mouse, we retained only the lineages that i) exhibited non-zero frequency over at least 12 out of 18 time points for the rm cohort and ii) the mean frequency over the entire time series is >=5e-5. Similarly, for the gf cohort which had 1 time-point less, we retained barcodes with i) non-zero frequency for at least 11 out of 17 time-points and ii) the mean frequency over the entire time-series is >=5e-5. For the im cohort, we keep lineages with i) at least 5 time points and ii) the mean frequency over the entire time-series is >=5e-6. This ensured that all barcode lineages included in the clustering had a sufficient number of points for pairwise comparison. This procedure meant that the lineage clustering focused on dominant and persistent clones; barcodes that immediately went to extinction were excluded. Altogether, this procedure was performed on a subset of ~300 to ~1300 lineages for each mouse, representing ~5% to ~10% of total barcodes. Since this analysis focuses on the dominant and persistent lineages, this fraction also represents ~7% to ~50% of the total number of *E. coli* cells (or raw barcode counts) at the end of the colonization experiment. The distance Δ*F_ij_* between two frequency trajectories *f_i_* an *f_j_* was calculated as

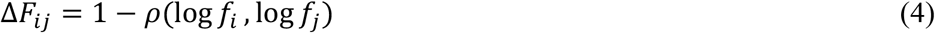

where *ρ*(log*f_i_*, log*f_j_*) is the Pearson correlation coefficient between the trajectories. A distance close to 0 indicated a strong positive correlation between the lineages, whereas a distance close to 2 indicated a strong negative correlation. From the resulting pairwise distance matrix, we applied hierarchical clustering using the “linkage” method from the *scipy.cluster.hierarchy* module in SciPy. We used the “average” agglomerative clustering method, which implements the algorithm unweighted pair group method with arithmetic mean (UPGMA)^51^. This method computes the distance between two clusters as the arithmetic mean of the distances between all lineages in both clusters. Then, for each cluster, we fitted a consensus trajectory using the local regression (loess). Loess is a form of moving average where a line is fit locally using neighboring points weighted by their distance from the current point. These moving averages were referred in the text as “clonal lineage clusters” or simply “clones”.

To determine the optimal clustering threshold, we note two general trends (Extended Data Fig. 2b-d). First, the loess of clusters with very few lineages will be sensitive to sequencing error. Thus, we include only clusters with at least 8 barcodes for the rm and gf cohorts and at least 5 barcodes for the im cohort. Second, when the threshold is too small, there are many clusters, but multiple clusters are similar to each other. This is manifested by the value of the smallest distance between the loess average of any cluster pair (black dots). Third, when the threshold is too large, there are very few clusters where barcodes with distinct dynamics are grouped together. In clustering, the practice was to find the cross-over between the smallest distance between cluster centroids (our loess average) and the number of clusters. This cut-off was indicated as the red curve in Extended Data Fig. 2b-d). Based on these cut-offs, we arrived at 4 to 21 clusters for im, 10 clusters for the rm cohort, and 6 or 7 for gf (Extended Data Fig. 3).

#### (v) Quantification of community dynamics by 16S profiling

The paired-end MiSeq Illumina reads resulting from sequencing of the 16S rRNA V4 region were processed using the *dada2* v1.22 pipeline^52^. Primer sequences were removed using *cutadapt* v2.8^53^ before amplicon sequence variant (ASV) inference. Forward and reverse read pairs were trimmed to a run-specific length defined by a minimum quality score (Phred score>= 25) using the *filterAndTrim* function of the *dada2* R package^52^. Error rates were estimated from sequence composition and quality by applying a core denoising algorithm for each sequencing run. Then pairs were merged if they overlapped using the *mergePairs* function. Bimeras, which were chimeric sequences, were removed with the *removeBimeraDenovo*. Taxonomy was assigned using the *assignTaxonomy* function that maps reads onto the *SILVA* (v. 138) reference database^54^. We excluded sequences that matched mitochondrial or chloroplast DNAs. In each mouse, the relative abundance of a taxonomic unit *i* at time *t* is given by:

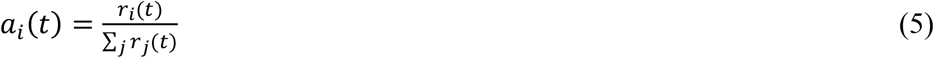

where *r*(*t*) is the absolute abundance (number of reads) for the unit. Similar to the barcode dynamics, we calculated the community’s effective diversity index but at the level of the family (see ***Quantification of barcode diversity***). For further analyses, families with frequency lower than 1e-03 were grouped as “Other”, while the rest of the groups were clustered under their bacterial family classification.

#### (vi) Co-clustering of *E. coli* clonal lineages and community dynamics from 16S

To detect the potential interactions between the bacterial community and *E. coli* clones, as might be manifested in the correlation between their time-series, we recognized that the interactions could introduce local and transient stretching or lags. Thus, a straightforward Pearson correlation is ill-suited to detect such interactions. Therefore, we calculated the pairwise distances using the shape-based metric (SBD)^36^. Briefly, the SBD is an iterative algorithm that detects the shape similarity of two time-series, regardless of amplitude or phase differences (Extended Data Fig. 5). For the community dynamics, we used the log-transformed relative abundances of taxa at the family level with a minimum of 7 non-zero time points. For the clonal dynamics, we used loess smoothing arising from the clustering of *E. coli* barcodes. We z-normalized the time series vectors to remove the amplitude effect and then calculated the shape-based distance (SBD)^36^ implemented in the *tsclust* package^55^ to calculate our distance matrix. Lastly, tree linkage was performed using the “average” (UPGMA) method to generate dendrograms (Fig. 2c and Extended Data Fig. 4c).

#### (vii) Assessing the biological significance of the co-clustering of clones and community dynamics

To validate that our co-clustering method between the community and clonal dynamics is significant, we calculated a metric called “mixing index”. The underlying rationale was that if indeed, clustering of an *E. coli* clonal lineage with a bacterial family is biologically meaningful, then this clustering should be strongest when both clonal lineage dynamics and 16S come from the same mice or same cohort. To assess the mixing index, we collect clone-clone *cophenetic* distances (*c*) and clone-species *cophenetic* distances (*m*) from their respective co-clustering. (Cophenetic distance is the distance between two leaves of a hierarchical tree and is defined as the height of the closest node that leads to both leaves). Then the distance between the empirical cumulative distributions of *c* and *m*, denoted as *F(c*) and *F*(*m*) respectively, is quantified as

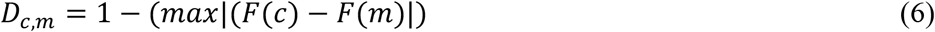

Higher values of the mixing index imply that clones and families are more likely to be adjacent leaves in the co-clustering three than clones amongst themselves. As an illustration, we show in Extended Data Fig. 5c the mixing indices for trees where clones and families are fully mixed, partly mixed, and fully unmixed. We applied the mixing index to co-clustering trees arising from different pairs of clonal lineages (im or rm or gf) and bacterial families (im or rm or nc). Furthermore, to determine the robustness of the mixing with respect to the method for determining the dominant clonal lineages (section iv), we evaluated the mixing index different cut-off thresholds for lineage clustering (Extended Data Fig. 2). The mixing index values are shown as violin plots in Fig. 3c-d. We found that the mixing index is largest when the clonal lineages and bacterial families come from the same mouse cohort. The statistical significance between the mixing indexes was quantified by a two-tailed t-test.

#### (viii) Replicability of clonal lineages in different mice from the same cohort

To determine the replicability of clonal lineage dynamics across different mice, we applied hierarchical clustering using distance matrices derived from pairwise Pearson correlation followed by UPGMA linkage (Fig. 5a,d). The input to these analyses was the loess average of the clonal lineages from each mouse (section iv. Barcode lineage clustering).

#### (ix) Quantification of barcode similarity between mice from the same cohort

To determine if the similarity in clonal lineage dynamics in different mice is driven by the same barcodes, we evaluated the overlap index in raw barcode identity for each cluster. In general, the overlap coefficient quantifies the Simpson similarity between two sets *A* and *B* that are not necessarily of the same size:

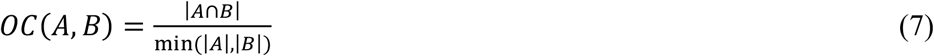

A value close to 1 indicates a high number of common elements, whereas a value near 0 indicates little overlap. We calculated the overlap index for all pairs of clonal lineage clusters in mice from the same cohort (see Fig. 5b,e). To determine that the overlap index did not arise by chance, we generated different compositions of sets *A* and *B* drawn randomly from our total pool of barcodes. For each composition, we calculated the overlap index (Eq. 7). This was performed 1000 times to arrive at a distribution of *OC*(*A,B*) values. The significance of the observed overlap index *x* between the real clusters *A* and *B* was expressed as a z-score on the simulated distribution of overlap indices:

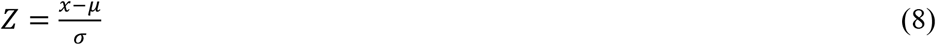

where *μ* is the mean and *σ* the standard deviation of the sample distribution. Lastly, significant overlap coefficient values with |*Z*| > 1.96 or p-value 0.05 are shown in blue in Fig. 6c,g, and their size is scaled proportionally to their p-value.

#### (x) Dynamic Covariance Mapping (DCM)

Microbes in species-rich communities participate in dynamic interaction networks whose time dependence can generally be quantified as a dynamical system by a set of ODEs (ordinary differential equations)^56^. However, a major limitation of existing methods for assessing the stability of nonlinear dynamical systems of ODEs is parametrization since they are not known a priori, and at best, they are inferred from noisy, sparsely sampled data. Here we developed a parameter-free methodology to quantify the Eco-Evo feedback of interactions on community dynamics from non-equilibrial time-series data. The dynamic covariance method (DCM) uses our unique high-resolution temporal data to quantify time-dependent interactions as they occur during the experiment. We start with the general case of a community composed of *N* members, representing *E. coli* clonal lineages and family-level bacterial taxa. A community vector *z*(*t*) can be defined as an *N*-dimensional vector of log_10_-transformed abundance time-series for the *E. coli* clonal lineages and family-level bacterial taxa:

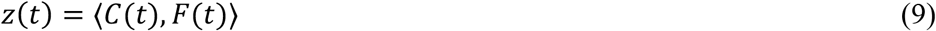

A non-detection in the community vector is replaced with a pseudo count of 1e-6. The vector *z(t*) describes the time-varying state of the community. Theoretically, its dynamics can be described by a system of ODEs

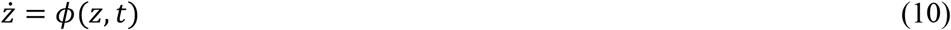

However, the functional form of *ϕ*(*z, t*) is unknown, but we can determine the generic behavior of the system near a specific snapshot (say, *z*_0_ = *z*(*t* = *t*_0_)) through linearization:

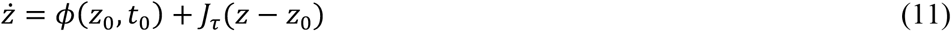

where *J_τ_* is the *N×N* Jacobian that defines the gradient of *ϕ*(*z, t*) approximated over the time interval *τ* around *t*_0_. The element of a Jacobian matrix measures the sensitivity of a species *i*’s population growth rate to the abundance change of species *j* and is defined as the interaction strength of species *j* on species *i* in an ecological community^57,58^; in practice, it can be estimated by the covariance of species *i*’ time derivatives and species *j*’s abundance time-series over the time interval *τ*:^9^

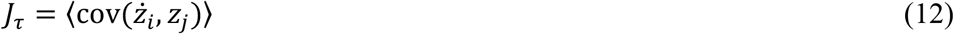

Here, subscripts *i* and *j* span from 1 to *N* and include both *E. coli* clonal lineages and family-level bacterial taxa. Following the dynamical systems theory^34^, the dynamics near (*z*_0_, *t*_0_) can be captured by the spectral distribution of eigenvalues (*λ_k_*, with *k* from 1 to *N*) of the Jacobian in the complex plane (representing both the real component Re(λ,) and the imaginary component Im(*λ_k_*)). The vector *z*(*t*) deviates from *z*_0_ at a rate of exp(Re(*λ_k_*)*t*) and oscillates at a period of 2*π*/Im(*λ_k_*) along the direction of the eigenvector associated with *Λ_k_*. The condition for *z*_0_ to be a stable equilibrium (i.e., the community can withstand small perturbations) thus requires Re(*λ_k_*) < 0 for all *k*.

In practice, we computed *J_τ_* over a specific time interval *τ* for progressively increasing time periods (3h-6h, 3h-12h, …, and 3h-15 days), with altogether a total of 16 or 17-time intervals for the rm and gf cohort and 7 or 8 17-time intervals for the im cohort. Since there are *N* eigenvalues for each time interval, we sought a simpler representation of the community’s dynamic behavior. To this end, we used Uniform Manifold Approximation and Projection (UMAP) ^59^ to reduce the dimensionality of the eigenspace as a function of time. The input to this UMAP dimensionality reduction is a (2*N*) × (4*n*) matrix, where the 2*N* columns correspond to the real and imaginary components of the eigenvalue *λ_k_* while the 4n rows correspond to the number of mice multiplied by the number of *τ* time intervals. In the UMAP’s 2D representation, the *2N*-dimensional eigenvalue is shown as a point (Fig. 4b, Extended Data Fig. 6e-g, Supplementary Movie 1a-d, and Supplementary Movie 2a-d). As shown in the Fig. 4b, the UMAP projection demonstrated that eigenvalues grouped together by time interval which suggests similar dynamic behaviors during colonization in different rm mice. Additionally, as the movie shows, the eigenvalues show distinct “jumps” on the UMAP projection, indicating distinct temporal phases. To define these distinct phases, we clustered the eigenvalues on their UMAP projection using the Nbclust^60^ package in *R*, implementing the *centroid* algorithm. To determine the robustness of identifying the distinct phases, we regenerated UMAPs using all possible neighborhood parameters (from 2 to 56, with the maximum value corresponding to the total number of points on the UMAP, i.e., 4n). We also tested other clustering algorithms (ward.D, ward.D2, single, complete, average, mcquitty, median, centroid, kmeans), which all showed 2 or 3 clusters (Extended Data Fig. 6a-d). In the case of the two clusters for the rm cohort, the first corresponds to *E. colf’s* entry and the second to its coexistence with the bacterial community. In the case of three clusters, there is an intermediate phase (the 2^nd^ phase) showing the resurgence of some bacterial species in the microbiota for the rm cohort.

## Acknowledgments

The authors thank Stephen Michnick and James Omichinski for discussions and Jesse Shapiro lab for protocols on 16S rRNA profiling. We thank the mice facilities at Université Sherbrooke and the University of Toronto. This work was supported by Canadian Institute for Health Research grant 408523 (AS), Canada Research Chairs (AS), National Research Foundation (South Africa) grant 89967 (CH), and a fellowship from Fonds de recherche du Québec - Nature et technologies (LG).

## Author contributions

AS, MG, and SB conceptualized this study. GMC, AF, and SR performed the E. coli colonization experiments in mice with innate and antibiotic-perturbed microbiota. DT and DP performed the colonization experiments in germ-free mice. MG performed the genomic extraction, barcode amplification, 16S rRNA profiling with the help of ZS. MG implemented all the bioinformatic analysis with the help of LG. AS, MG, and CH developed the dynamic covariance mapping approach. AS and MG wrote the the manuscript with feedback and editorial support from all authors.

---

**Extended Data Fig. 1.**
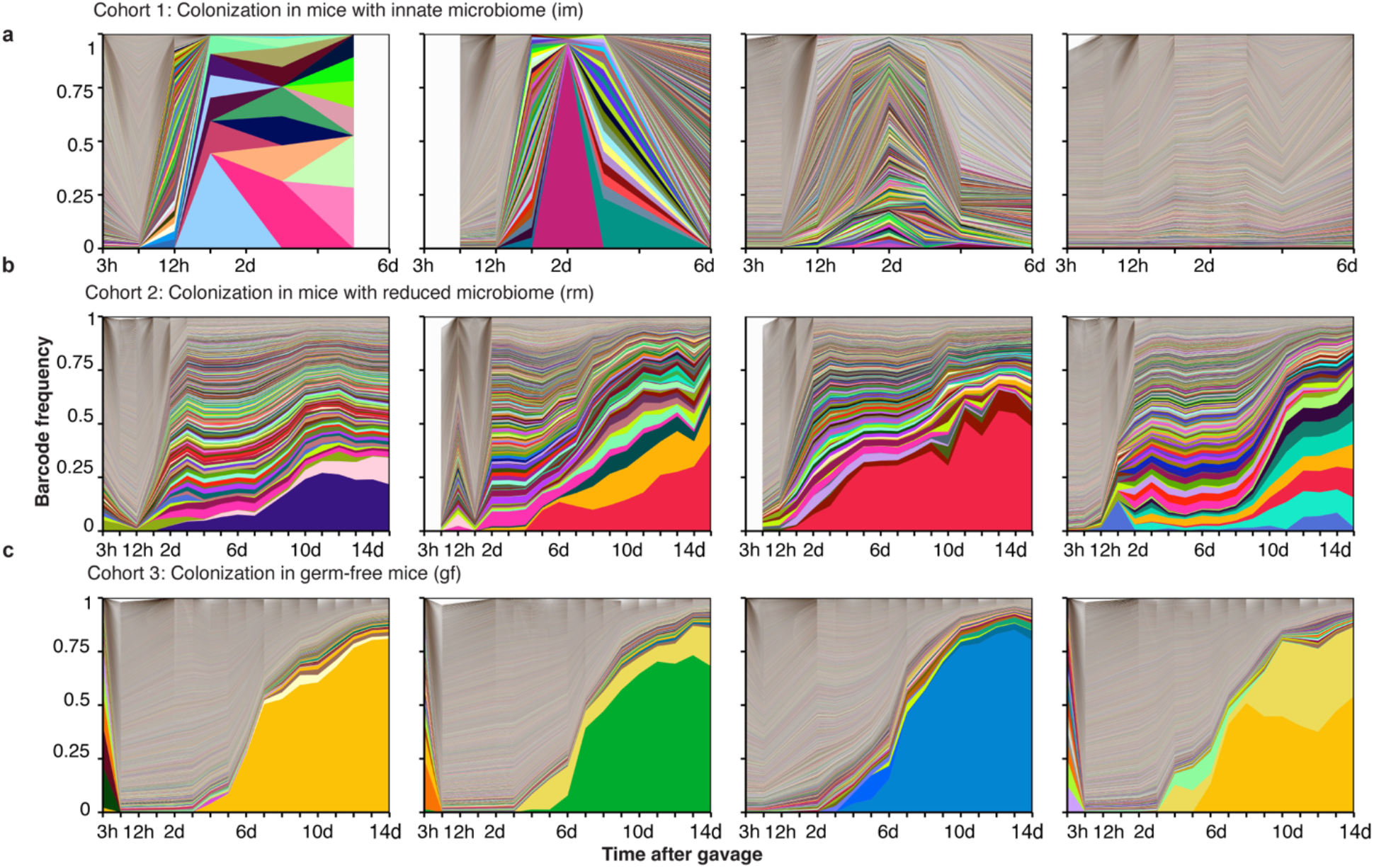
Barcode dynamics in im, rm, and gf cohorts. **a-c,** Barcode dynamics for cohort 1, cohort 2, and cohort 3 in linear scale. Each column corresponds to replicate mouse 1 to 4, respectively. The color corresponds to Fig. 1c-e.

**Extended Data Fig. 2.**
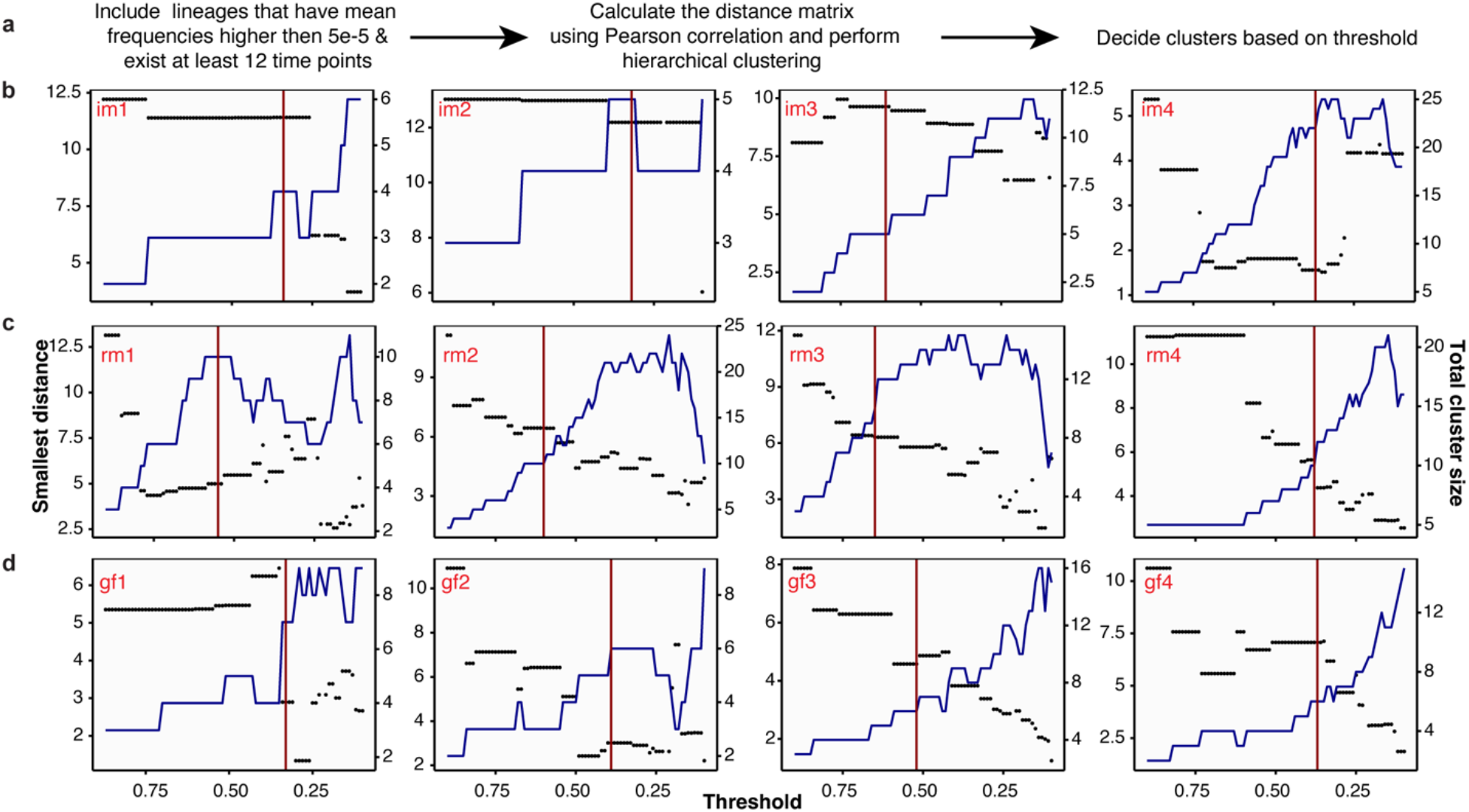
Determining the number of dominant clonal lineages. **a,** The pipeline for estimation of putative clones from the frequency time series of the chromosomal barcodes. **b-d,** A specific value for the threshold distance (Pearson correlation) in the hierarchical clustering defines a total number of clusters (blue curve) as well as a distance between the most similar clones (“Smallest distance”, black dots). When the threshold is large, there are many clusters, but some are similar to each other. Conversely, when the threshold is small, there are too few clusters, where even barcodes that do not have similar time series are grouped together (Methods). In practice, the cut-off is chosen to be the cross-over between the smallest distance between cluster centroids (our loess average) and the number of clusters. The chosen cut-off is indicated by the red curve. The resulting clusters are shown in Extended Data Fig. 3.

**Extended Data Fig. 3.**
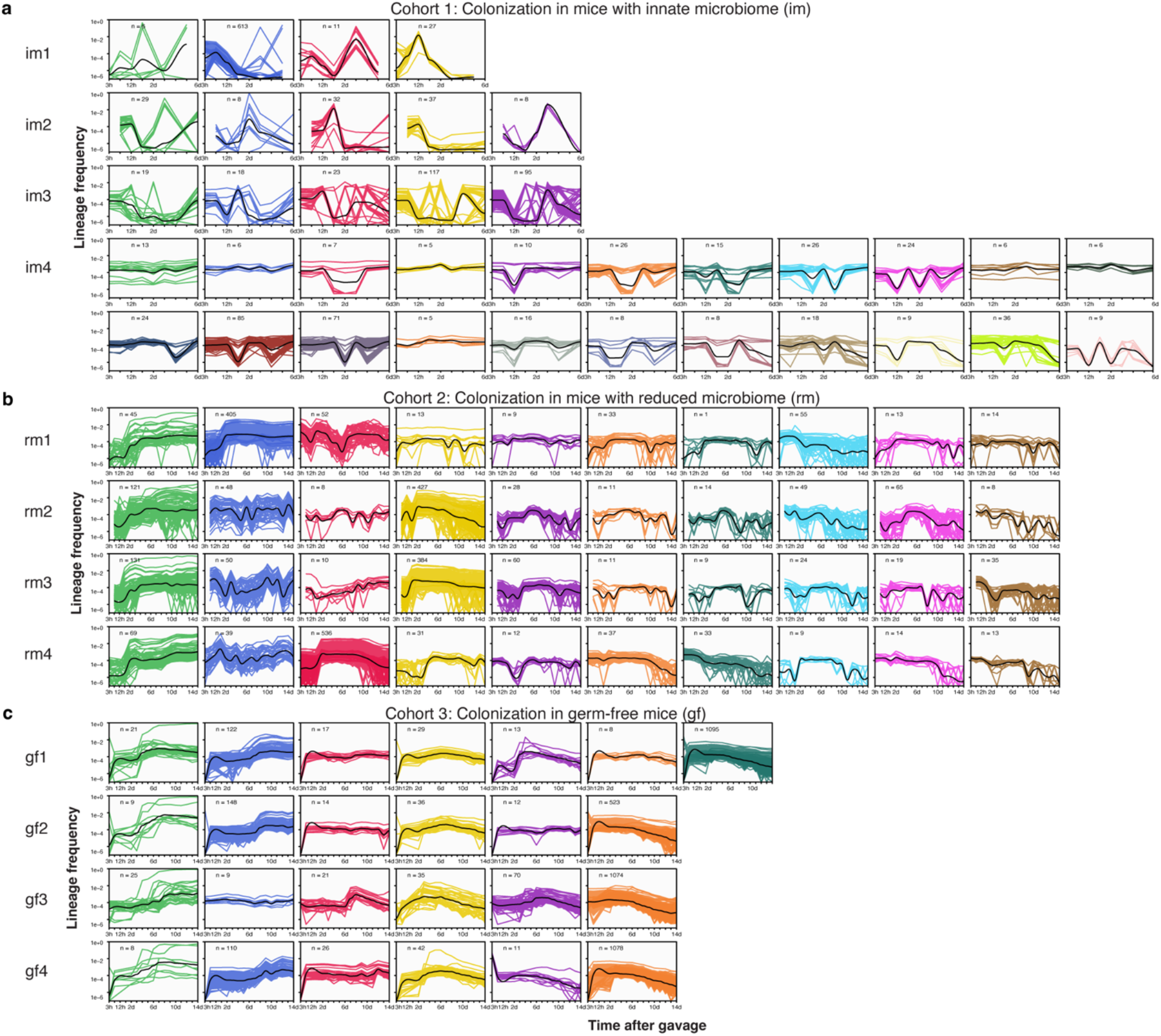
Dominant E. coli clonal lineages clusters. **a,** Dominant barcode clusters in mice with innate microbiota (im). Colored lines correspond to unique chromosomal barcodes in the cluster. Black lines correspond to the LOESS average. The number of unique raw barcodes that belong to the cluster is indicated. The clonal lineage clusters (or simply “clones”) are ordered, starting from the left, based on their average barcode frequency on the last day. **b**, Dominant clusters for the mice with reduced microbiota (rm). The colors correspond to Figure 2a. **c**, Dominant clusters for the germ-free mice (gf). The colors correspond to Figure 3a.

**Extended Data Fig. 4.**
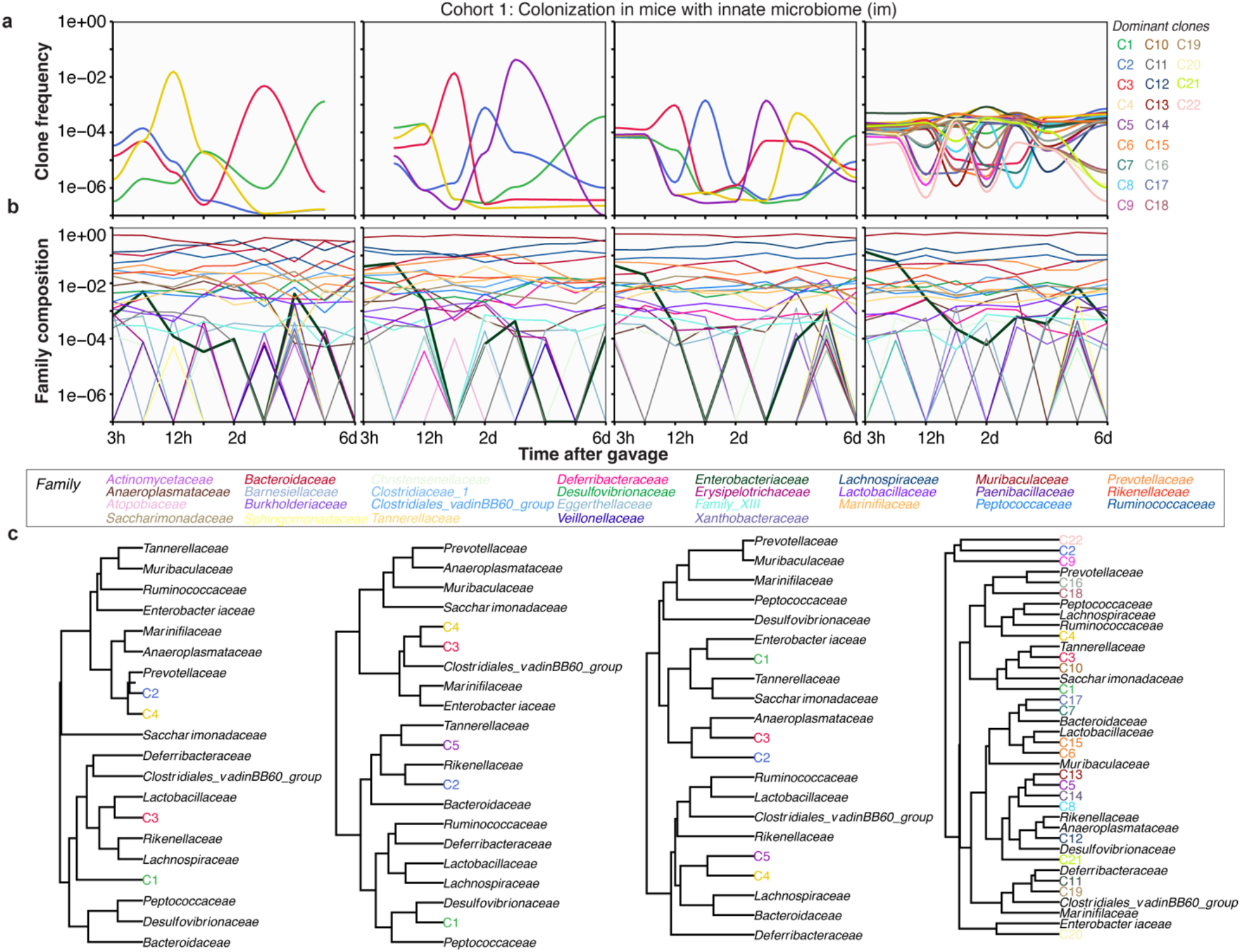
Clone-specific community interactions in the gut for the im cohort. **a,** Dominant and persistent *E. coli* clones invading an innate microbiome show less reproducible clonal dynamics compared to gf and rm cohorts. Particularly, mice im1-3 have less than 6 clones, but im4 has 21 distinct clones. **b,** Community dynamics are similar for im1-3 where the *Enterobacteriaceae* (thick line), to which *E. coli* belongs, drops below the resolution limit of the 16S rRNA profiling but persists in im4. Interestingly, this distinction in dynamics between mice im1-3 and mouse im4 is not perceptible from the CFU (Fig. 1b).

**Extended Data Fig. 5.**
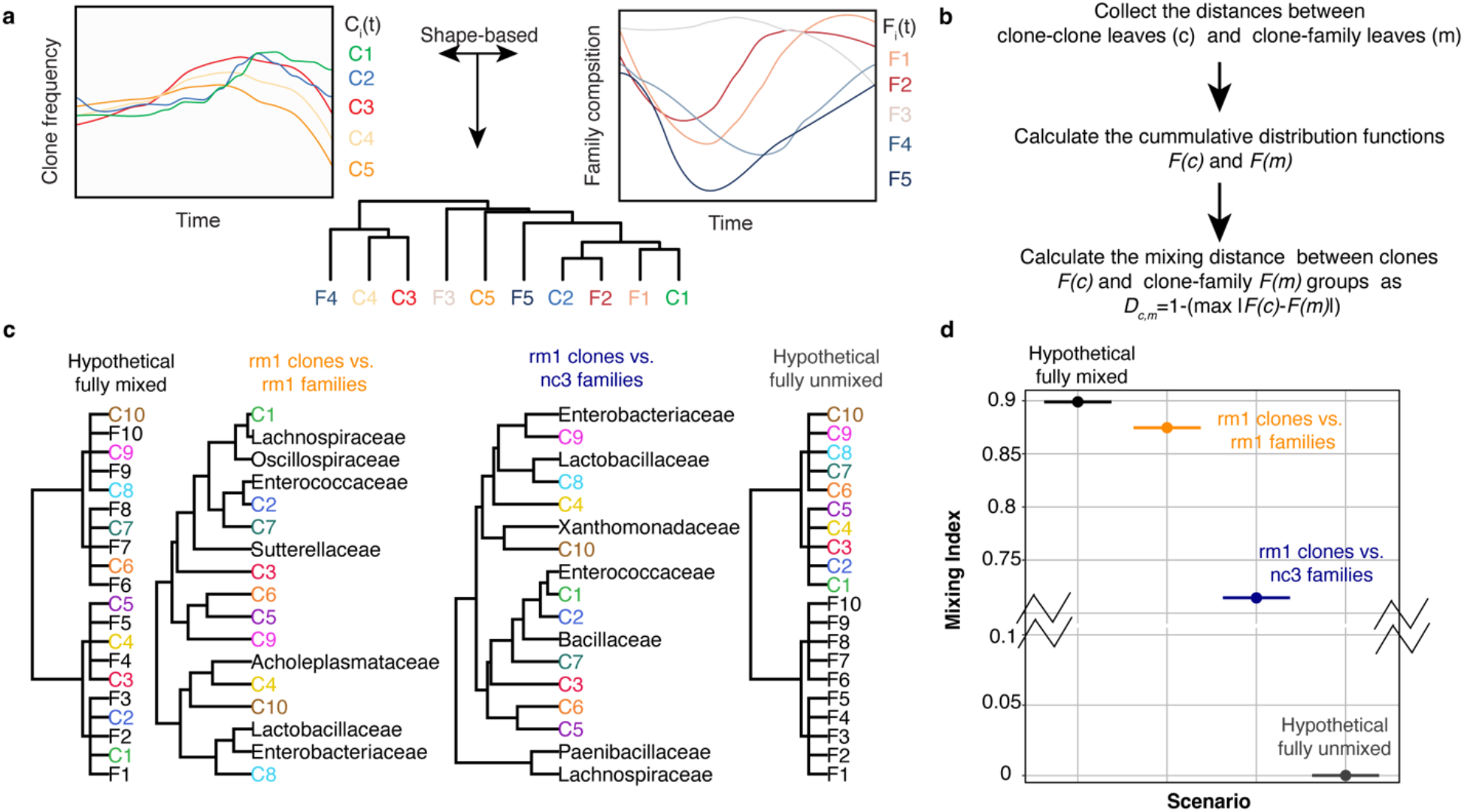
Schema for shaped-based co-clustering between E. coli clone and community dynamics. **a,** Schema for co-clustering of the clonal lineage and family composition time series using a shape-based distance metric. **b**, Schema for calculating the mixing index from a given hierarchical clustering tree. We collect clone-clone *cophenetic* distances (c) and clone-species *cophenetic* distances (*m*). Then the distance between the empirical cumulative distributions of *c* and *m*, denoted as *F*(*c*) and *F*(*m*) respectively, is quantified as *D_cm_* = 1 – (*max*|(*F*(*c*) – *F*(*m*)|). Higher values of the mixing index imply that clones and families are more likely to be adjacent leaves in the co-clustering tree than clones amongst themselves. **c-d**, Illustrative examples (panel c) of different extents of co-clustering between clones and families and their corresponding mixing indices (panel d).

**Extended Data Fig. 6.**
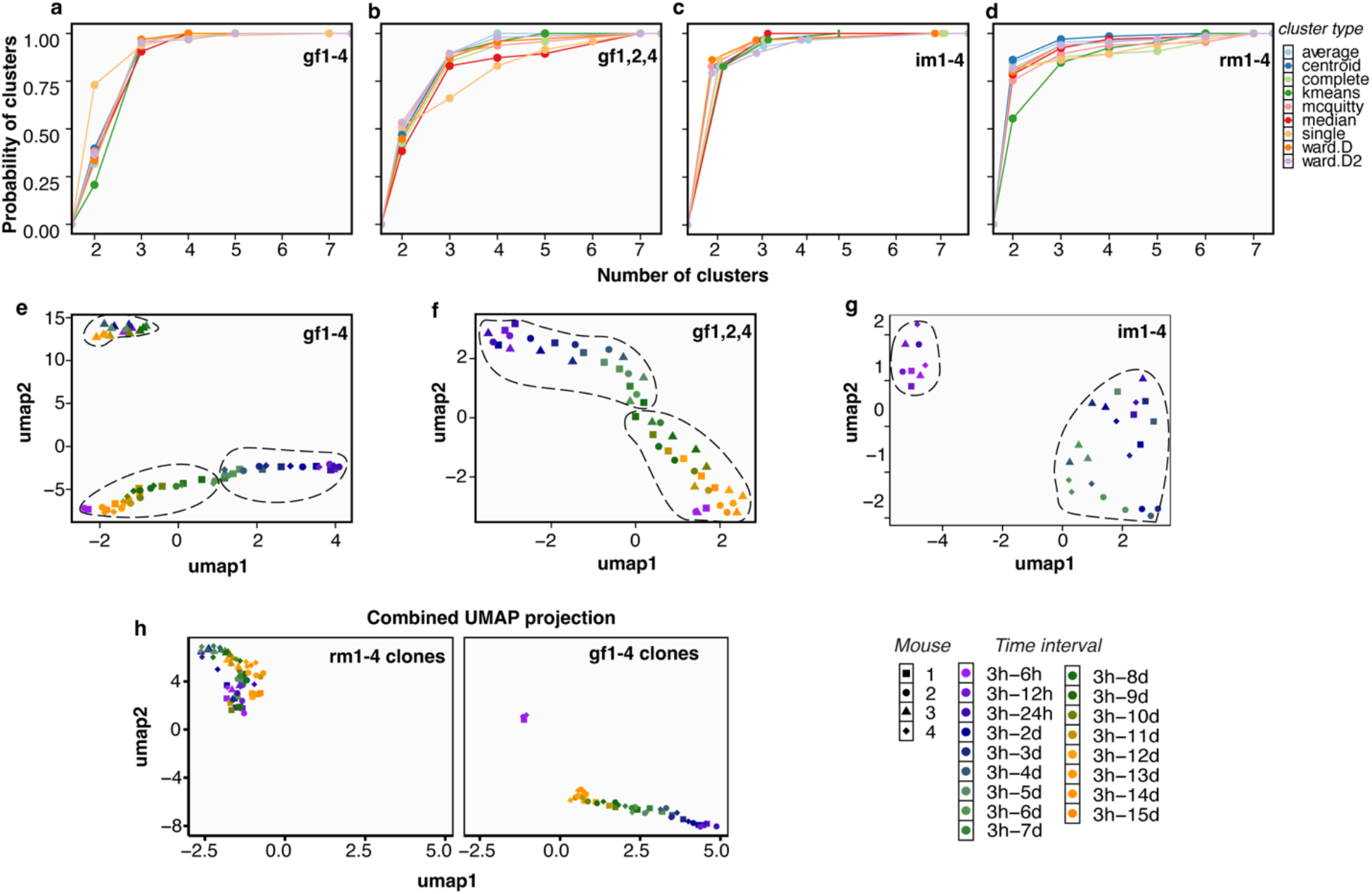
Robustness of UMAP clustering and DCM analysis. **a,** We generated all possible UMAPs for the eigenvalues of the gf cohort using every possible value of the n-neighbor criteria (Methods and ref. ^59^). Then, we applied nine different clustering algorithms to determine the groupings of eigenvalues on these UMAPs. These groups indicate the phases of invasion defined in the text. Shown is the probability of the number of groups (clusters). Robustly, there are 2 or 3 groups or phases on the UMAP. **b**, Similar to panel a, panel b shows that the presence of 3 clusters in the four gf mice is driven by gf3. **c-d,** Similar to panel a for the invasion in mice with innate (**c**) and reduced microbiota (**d**), respectively. **e**, The chosen UMAP for the gf cohort (panel **a**). **f**, The gf cohort, where gf3 is excluded, shows distinctly 2 phases (see also Supplementary Movie 2a-d). **g**, The phases defined on the UMAP of the im cohort. **h,** We combined all eigenvalues from the rm clone dynamics and gf cohort and projected them onto a single UMAP. The rm clones are shown on the left panel, while the gf clones are shown on the right. Projection of rm and gf cohorts’ clonal lineages on the same UMAP shows that rm1-4 clones grouped amongst themselves together with gf3. At 3h, gf3 already exhibits high barcode diversity, indicating that most of the barcode has transited the gut (Fig. 1f, 3a third panel, and Extended Data Fig. 1c third panel), in contrast to the other mice in the gf cohort.

**Extended Data Fig. 7.**
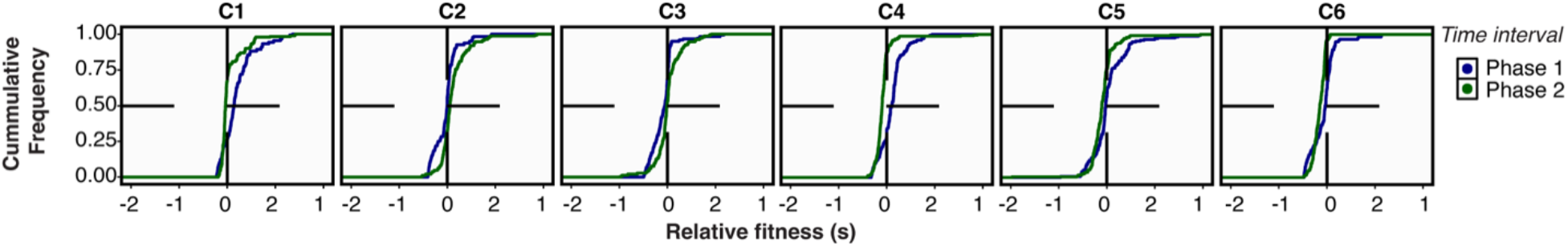
Estimated fitness of the gf cohort. The cumulative distribution is partitioned into the two phases defined in Extended Data Fig. 6f. The dominant clone C1 exhibit primarily adaptive dynamics during phase 1 of the colonization.

## Notes

### Competing Interest Statement

The authors have declared no competing interest.

